# Heterologous engineering of receptors using OrthoRep (HERO) for directed evolution of GPCRs in yeast

**DOI:** 10.64898/2026.09.30.755785

**Authors:** Christien B. Dykstra, Bjorn D.M. Bean, Olivier Rousseau, Flavia Araujo, Dima Masoud, Chang C. Liu, Malcolm Whiteway, Vincent J.J. Martin

## Abstract

Engineering G protein-coupled receptors (GPCRs) for biosensing applications remains challenging due to structural and functional constraints when such proteins are expressed in a heterologous host. OrthoRep offers the continuous accumulation and selection of signal-enhancing mutations in yeast, yet it has not been applied to GPCR evolution. We apply OrthoRep-driven mutagenesis to evolve opioid GPCRs and improve yeast-based biosensors: first, we express the human μ-opioid receptor (OPRM1) on the OrthoRep p1 plasmid and couple ligand activation to growth. Serial cell passaging of the biosensor generated functionally diverse mutants. However, mutations that decouple receptor activation from selection arose over longer passaging campaigns, halting evolution. We resolve this issue using a site-specific recombinase that re-introduces evolved variants onto the p1 landing pad of an unmutated biosensor, allowing continued selection. This led to an ∼18-fold improvement in sensitivity of a mutant OPRM1 over wt-OPRM1. Additional polymerases were added to the platform, broadening the mutational profiles available. Assembling these components forms the platform we call HERO (Heterologous Engineering of Receptors using OrthoRep). Lastly, we deploy HERO using automation to improve the human δ-opioid receptor’s weak response to an agonist by ∼40-fold. Sensitivity also improved for a structurally dissimilar ligand assayed on the same evolved variants.

## INTRODUCTION

Although natural selection has produced sequence solutions that elude those of current rational design, it’s a path-dependent process that retains only those modifications accessible by fitness-neutral or - improving mutational steps from a pre-existing template. As a consequence, continuous directed evolution is an attractive route to engineering novel protein function precisely because it escapes those paths: the experimenter defines the selection pressures, starting points, and mutational regimes^1^. One such continuous approach is OrthoRep^2^ which is an orthogonal DNA replication system in the yeast *Saccharomyces cerevisiae* adapted from the linear, cytosolic p1/p2 killer plasmids of *Kluyveromyces lactis*^3^. p1 encodes a killer toxin and its own terminal-protein-bound DNA polymerase (TP-DNAP1), and p2 encodes a second p2-targeted polymerase termed TP-DNAP2, along with the RNA polymerase and housekeeping genes needed to maintain both plasmids. Only the translation machinery is supplied by the host (Supplementary Fig. 1A). The OrthoRep system replaces most of p1’s payload with a gene of interest (GOI), and expresses an engineered, mutagenic TP-DNAP1 *in trans*. The GOI continuously accumulates mutations during replication, at rates up to 10^-4^ substitutions per base (s.p.b.), while selection acts on the resulting variants in real-time^2,4,5^. Because OrthoRep was developed for yeast, a eukaryotic host, it provides the post-translational processing and signaling context that receptors require to function. Thus, continuous evolution of protein classes defined by these dependencies is a worthwhile possibility. However, unlike enzymes, receptors generally report function indirectly through multi-step signalling cascades. Keeping that coupling faithful across many generations of mutagenesis and selection rounds presents a challenge, and receptors have therefore remained an under-exploited target for this approach.

G-protein coupled receptors (GPCRs) are seven-transmembrane domain proteins, ubiquitous across eukaryotes, that detect stimuli ranging from pheromones to light^6^. Ligand binding triggers a conformational change that is transmitted across the membrane, allowing GPCRs to engage intracellular G proteins and elicit downstream responses. In humans, this core signalling architecture has been repeatedly repurposed, giving rise to over 800 GPCR members across five classes, an evolvability that also makes them attractive substrates for directed evolution toward new sensing functions^7-9^. They are also the target of ∼36% of all approved drugs, reflecting their importance in therapeutic screening^10^. Medically important examples are the μ-, κ-, and δ-opioid receptors (OPRM1, OPRK1, OPRD1), which couple to inhibitory G_i_/G_o_ proteins, lowering cAMP to produce the analgesic effects of widely used pain medications. Hundreds of structurally diverse ligands have been characterized at OPRM1, and libraries of thousands of compounds have been screened for therapeutic potential^11^, yet functional screening of these compounds in mammalian systems is comparatively costly and low-throughput. Indeed, yeast-based biosensors have already been put to pharmaceutically relevant use, from screening ligands such as cannabinoids^12^ to point-of-care detection of fungal pathogens^13,14^; however their widespread deployment is limited by the ability to functionally express targets for compound screening.

We previously built functional opioid biosensors by coupling human opioid receptors to yeast’s native pheromone response pathway, reading out activation through a fluorescent reporter^15^. Replacing ergosterol with cholesterol in these strains improved the opioid receptors’ sensitivities in the yeast biosensor context, but they remained considerably lower than their reported responses in mammalian cells. Additionally, the resulting strains grew poorly and were difficult to transform, limiting their practicality as a screening platform. Receptor-level engineering offers a complementary route to more sensitive biosensors, one that OrthoRep is well suited to pursue by continuously evolving GPCRs such as OPRM1 directly in the yeast host. Biosensors of this kind could be used for pathway engineering capable of detecting improved metabolite production by enzyme variants, perhaps under continuous evolution themselves. Because GPCR sensing is extracellular, these biosensors are also suited to reporting impermeant and complex metabolites or peptides that other intracellular biosensor configurations cannot^9,16,17^. Beyond expanding the opioid biosensor toolkit, receptor evolution could begin to reach more distant regions of GPCR sequence space, in principle neofunctionalizing receptors toward ligands they do not natively recognize. Even where the target ligand is itself well known, such as a benzylisoquinoline-alkaloid intermediate like reticuline, a receptor that detects it would constitute a new and useful sensing function^18^.

With the goal of expanding our opioid detection platform, we successfully demonstrate continuous, directed evolution of opioid receptors in yeast: we have established an OrthoRep biosensor that couples receptor activation to yeast growth, which we develop across three phases. In the first phase, we show that passaging biosensor cells in the presence of saturating ligand generates functionally diverse OPRM1 variants that are tolerated by the host and read out across different dynamic ranges, but that this diversification alone does not reliably yield variants more sensitive than wild type. We also anticipated that coupling mutagenesis to a reporter several steps downstream of the receptor would introduce a recurring failure mode, selection escape, because mutations in the MAPK pathway downstream of receptor activation can drive ligand-independent activation of the growth reporter^19-24^. To resolve this, in the second phase we implemented a refreshable landing-pad system in which evolved variants are amplified from OrthoRep and rapidly re-installed onto p1 in an unmutated biosensor background using recombinase-mediated integration^25^. This leaves behind the biosensor chassis that escaped selection, and carries the receptor population forward for continued rounds of evolution on OPRM1^26^. Assembling these components yields the platform we call HERO (Heterologous Engineering of Receptors using OrthoRep), a plug-and-play system for continuous GPCR evolution and screening. Having shown that HERO, seeded with evolved sequences from the earlier phases, increases OPRM1 sensitivity to a peptide opioid, we demonstrate HERO’s utility in the third phase by markedly increasing the sensitivity of yeast expressed OPRD1 to a synthetic opioid 40-fold over wild type using automation-assisted cell passaging.

## RESULTS

### Adapting OrthoRep for mutagenesis of an opioid receptor within a yeast-based biosensor

We built an OrthoRep growth-based yeast biosensor that only grows when its target ligand is present, reasoning that with repeated passaging, receptor mutations that improve agonism and signal transduction would be enriched (Fig. 1A). Our biosensor chassis is adapted directly from Bean *et al*.^15^, where the yeast pheromone-response pathway was rewired so that mammalian opioid receptors induce the expression of a fluorescent reporter. In this previously characterized biosensor chassis, the native GPCR, *STE2*, was deleted to eliminate interference at the top of the pathway^27-29^, and the final five residues of Gpa1 (K468IGII → ECGLY) were modified to have the yeast Gα mimic the human opioid partner Gαi3^30^ (referred to as Gαi3 hereon). *FAR1* and *SST2* were also deleted to stop agonist-induced cell-cycle arrest and to relieve signal shut-off, respectively (Fig. 1B)^31,32^. As the cholesterol-producing versions of this chassis displayed reduced growth, we only adapted the ergosterol strain. In addition, we replaced the original fluorescent reporter with *HIS3*, which encodes a histidine biosynthesis enzyme. In our *his3Δ* background, cells grow without histidine only when the pheromone pathway drives *pFIG1-HIS3*, allowing growth as a measure of receptor activation^15,33^. Further, histidine production by *HIS3* can be competitively inhibited by 3-aminotriazole (3-AT), allowing tunable selection and suppressing background from basal expression during future cell passaging (Fig. 1B)^34,35^.

**Figure 1.**
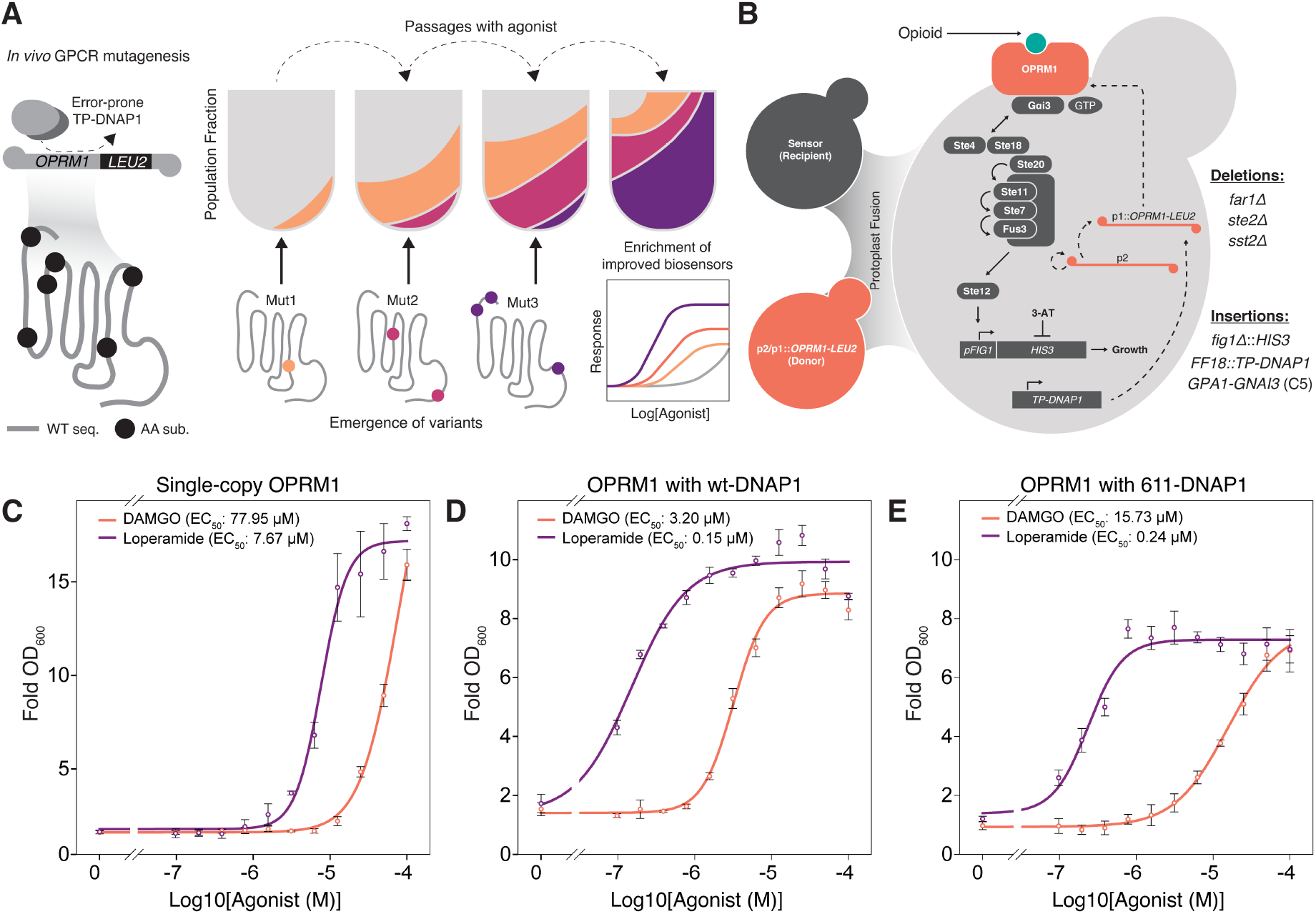
A growth-based OrthoRep biosensor functionally expresses OPRM1. **A** Overview of continuous directed evolution of opioid biosensors: an error-prone orthogonal polymerase (TP-DNAP1) mutagenizes the p1-encoded GPCR in a growth-based yeast biosensor. Populations are serially passaged into fresh medium, accumulating substitutions across generations, allowing the enrichment of mutations that improve signalling and growth over the wt/deleterious pool. The end results are biosensors with improved signalling over wt. **B** Schematic of the OrthoRep yeast biosensor assembled by protoplast fusion of a recipient (sensor) carrying *FAR1, STE2*, and *SST2* deletions together with *GPA1-GNAI3* (Gαi3), *pFIG1-HIS3*, and user-defined *TP-DNAP1* insertions with a donor supplying p2 and p1::*OPRM1-LEU2*. In this fusant, agonist-bound OPRM1 activates the Gαi3 chimera to signal through the cascade to a pFIG1-HIS3 reporter, while an error-prone TP-DNAP1 replicates p1. Further, growth may be tuned via the addition of 3-AT. Growth-based dose-responses using a Log_2_ gradient of **C** the single-copy OPRM1 control, **D** the wt-DNAP1 fusant expressing OPRM1, and **E** the mutator 611-DNAP1 fusant expressing OPRM1. Points are mean ± s.d.

Because both strains share the same mating type (*MAT*a), and because *Δste2* removes mating competence, protoplast fusion was chosen to introduce the cytosolic p1 and p2 components into our biosensor chassis, allowing us to repurpose our already extensively edited sensor chassis^2^. We fused a donor carrying modified p1::*OPRM1-LEU2* together with wild-type p1 and p2 to a recipient biosensor bearing a genomically integrated mutagenic DNAP1 (Fig. 1B, Supplementary Fig. 1). Selection on -Leu/-Trp required the recipient *TRP1* marker and the donor-borne *LEU2*, respectively, so that only recipients that had acquired the donor’s cytoplasmic plasmids would be recovered. The intended product is a haploid recipient carrying its own sensor genome with only the transferred cytoplasmic p1 and p2 generating a functional OrthoRep biosensor. We tested three TP-DNAP1s from Ravikumar *et al*.: wt-DNAP1, 611-DNAP1, and DNAP1-4-2, with approximate error rates of 10^-9^, 10^-6^, and 10^-5^ substitutions per base (s.p.b.)^4^. The wt-DNAP1 rate is close to that of genomic replication, so this strain serves as a low-mutation control and a baseline sensing benchmark. wt-DNAP1 and 611-DNAP1 both produced appreciable amounts of fusants under - Leu/-Trp selection (Supplementary Fig. 2A). However, DNAP1-4-2 did not yield a functional biosensor after repeated attempts, potentially because of its high error rate/low processivity on p1/p2 during the post-fusion window preventing the recovery of a working receptor^5^. After protoplast fusion, transient multinucleation can arise, but progression to stable nuclear hybrids occurs in well under 1% of fused pairs^36^. Propidium iodide staining confirmed both wt-DNAP1 and 611-DNAP1 recipients retained haploid DNA content after one passage in -Leu/-Trp, while DNAP1-4-2 gave aberrant results (Supplementary Fig. 2B,C). Despite its lower mutagenic rate, we carried forward with 611-DNAP1 for our initial evolution campaigns.

To test OPRM1 evolution in our 611-DNAP1 strain, we picked two structurally distinct μ-opioid receptor agonists: DAMGO, a synthetic peptide, and loperamide, a small-molecule piperidine. Evolving against agonists from different chemical classes lets us test whether the platform behaves consistently across dissimilar ligands. To benchmark the growth-based opioid biosensor system, we first built a single-copy genomic wt-OPRM1 biosensor in the recipient chassis by placing OPRM1 under the strong *PGK1* promoter. This construct also served as an unmutated control to validate single-copy, growth-based screening of evolved variants during later runs. Growth was read at a 48-h endpoint, matching the density cultures required to reach levels acceptable for passaging the cells in the evolution campaigns. Loperamide gave an EC_50_ of 7.67 μM, while the DAMGO response did not saturate within this window despite a robust signal (fold change ∼16; Fig. 1C). Both fusant strains, by contrast, gave clean sigmoidal curves: the wt-DNAP1 strain reached EC_50_ values of 3.20 μM (DAMGO) and 0.15 μM (loperamide) (Fig. 1D), while the 611-DNAP1 strain was right-shifted (15.73 μM and 0.24 μM; Fig. 1E). The fusants express OPRM1 from the multi-copy p1 plasmid rather than a single genomic locus, so their greater sensitivity relative to the control is consistent with a receptor-dose effect^33^, and across the two fusants, sensitivity tracked p1 copy number, the higher-copy wt-DNAP1 strain being more sensitive than the lower-copy 611-DNAP1 strain^5^.

### Generation of host-tolerated functionally diverse opioid receptor variants

In the first phase of OrthoRep biosensor development, we ran cell passaging campaigns as four independent replicates of the 611-DNAP1 biosensor in 2 mL SC-His/Leu (pH 7.1), supplemented with either 50 μM DAMGO for ∼90 generations (14 cell passages; Fig. 2A) or 50 μM loperamide for ∼60 generations (10 cell passages; Fig. 2B). We used near-saturating agonist concentrations to keep selection pressure on the receptor low, and reasoned that lower-activity intermediates could persist rather than being purged, leading to a greater diversity of solutions rather than a rapid convergence on a few^37^. Independently, 3-AT competitively inhibits the His3 reporter, providing a second control that sets how much signal is required to grow and suppresses leaky, ligand-independent background growth. Titrating 3-AT against DAMGO in the wt-DNAP1 strain, growth at 2.5 mM rose with agonist concentration without saturating at 48 or 72 h (Supplementary Fig. 3), confirming that output still tracked signal strength; we used 2.5 mM in parallel campaigns alongside the primary condition. The loperamide campaign was initiated later and run for fewer passaging rounds; the two are therefore compared by evolutionary trajectory as a function of the resulting substitutions, rather than by substitutions per generation. For each round, agonist-treated cultures were back-diluted 1:100 into fresh media with agonist, alongside a control group that tracked each population’s tendency to escape selection. Cultures grew for 48 h before OD reading and the next passage. Parallel passages received 2.5 mM 3-AT to test whether raising *HIS3* selection stringency could alter the mutational profile in comparison to agonist only (Supplementary Fig. 4). The alternating high- and low-growth passages seen in all groups (Fig. 2A and B; Supplementary Fig. 4) most likely reflect the fixed 48-h passaging scheme rather than population instability: proliferative cultures reaching saturation sooner may comprise more unfavorable variants at the time of passaging as a function of cell count, diluting the fraction of improved variants. The next passage begins under effectively tighter competition, allowing improved variants to reclaim a greater share of the next starting population, and the cycle repeats.

**Figure 2.**
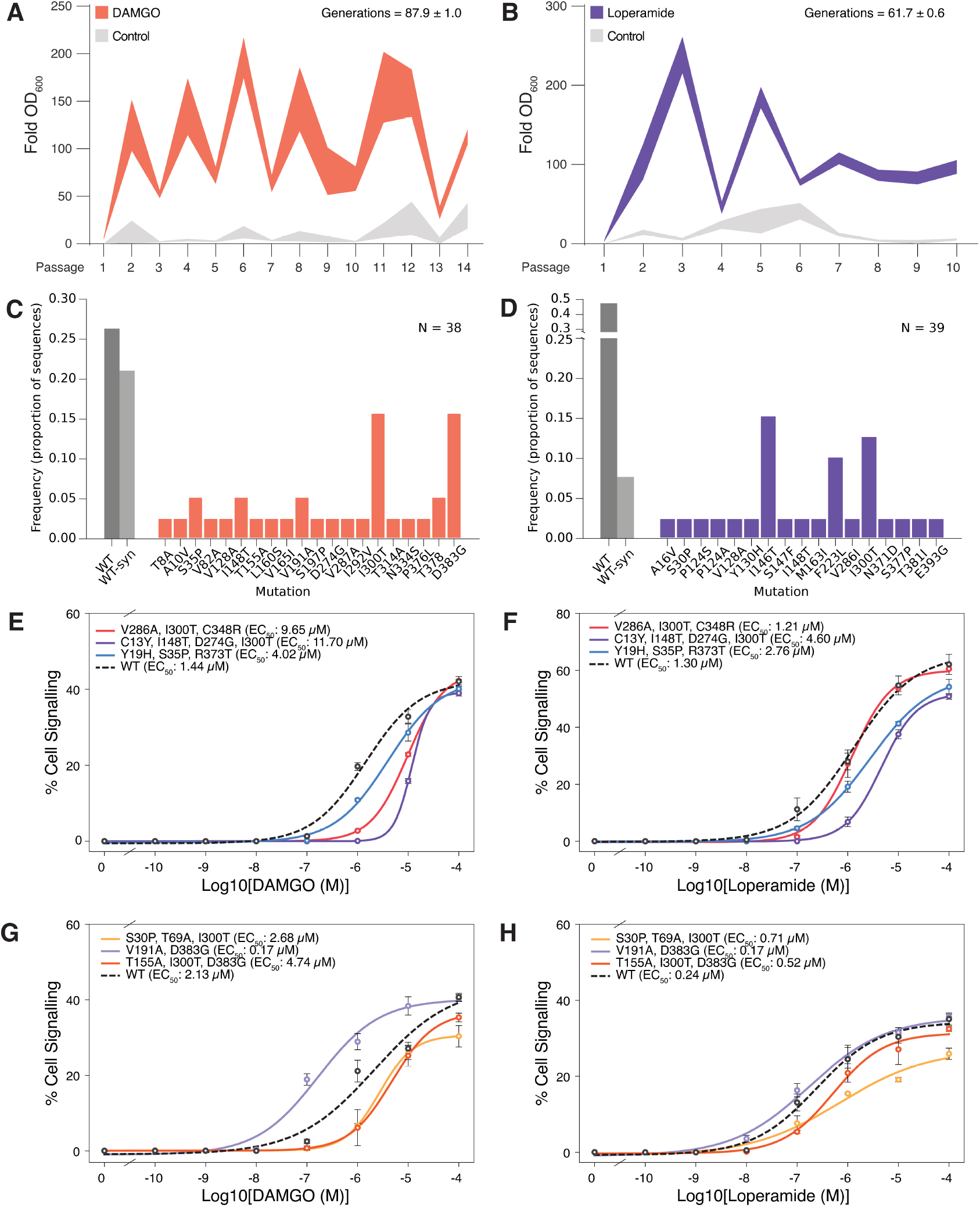
Passaging of biosensors with OrthoRep generates tolerated, functionally diverse OPRM1 variants. **A** Growth (OD_f_/OD_i_) of 611-DNAP1 biosensor expressing OPRM1 across 14 DAMGO passages and **B** 10 loperamide passages, each with an agonist and control groups (N = 4, fill is s.d.). **C** Endpoint substitution frequencies for the DAMGO-evolved, counter-selected, fluorescence-sorted pool (N = 38); wild-type and WT-synonymous classes are grey and nonsynonymous substitutions colored, with I300T and D383G most frequent. **D** Endpoint substitution frequencies for the loperamide pool (N = 39), which enriched I146T and F223L. **E** DAMGO and **F** loperamide dose-responses measured by flow cytometry of isolated DAMGO-selected variants (V286A/I300T/C348R; C13Y/I148T/D274G/I300T; Y19H/S35P/R373T) against a wild-type reference. **G** DAMGO and **H** loperamide dose-responses of a second variant panel measured by flow cytometry (S30P/T69A/I300T; V191A/D383G; T155A/I300T/D383G), in which V191A/D383G is markedly more sensitive than wild type. **E-H** use a Log_10_ gradient, points are mean ± s.d.

To screen single receptor variants from the evolved pools, the OPRM1 variant pool was recovered from p1 and integrated as a single genomic copy into the biosensor chassis carrying a *pFIG1*-driven fluorescent reporter (*ENVY*). This prevented further mutating of the receptor or confounding signalling gains and sequencing results with p1 copy-number or heteroplasmic background. We then sorted by fluorescence against the top 30% of wt-OPRM1 signallers (Supplementary Fig. 5A)^38^. The mutated OPRM1 population showed a decrease at this gate, indicating a pool of non-functional variants that may emerge at any point during the passaging campaign^39^. After ∼35 generations (passage-6), the DAMGO-selected library showed clear mutation accumulation, with means of 1.3 nucleotide and 0.8 amino-acid substitutions per sequence (Supplementary Fig. 6A,B).

The most frequent single substitution was I300T, present in 39% of sequences. A protein-level WT fraction of 0.46 (WT plus synonymous-WT) indicates that much of the enrichment favours wt-OPRM1 at this early stage of evolution, a fraction that declines over later passages. The mutation profiles of the ±3-AT groups did not differ at this point (Supplementary Fig. 6C). Endpoint sequencing showed that each selection condition drove its own mutational spectrum. In the DAMGO-evolved, fluorescence-sorted pool (n = 38), I300T and D383G were the most frequent substitutions, appearing alone or together, and in both DAMGO-only and 3-AT pooled populations (Fig. 2C). The loperamide pool (n = 39) also enriched I300T but additionally selected strongly for I146T and F223L (Fig. 2D). Aside from the shared I300T, these enriched substitutions recurred across independent sorted clones: D383G seen only in the DAMGO pools versus I146T and F223L seen only in the loperamide pool. Because the pools were processed separately until barcoding, the recurrence of I300T across these independent streams is most consistent with positive selection rather than neutral carryover from the starting strain. Its target of selection is less clear: I300T could be a permissive change acting on a general property such as folding, expression, or basal coupling, but its further enrichment under non-selective screening (below) suggests its advantage may lie partly in growth or propagation rather than ligand-driven signalling.

A heteroplasmic starting pool carrying both WT (favoured under agonist selection) and I300T (favoured during maintenance) could reconcile its early presence with ongoing selection, though disentangling these dynamics is beyond our present scope. After roughly 90 (DAMGO) and 60 (loperamide) generations, about half of the coding sequences remained WT or synonymous-WT, a higher WT fraction than the per-base error rate alone would suggest^4,39^. This likely reflects two non-exclusive dynamics: in some lineages the WT receptor still carries most of the productive signal and co-occurring mutations are neutral or mildly harmful hitchhikers carried along by clonal expansion; in others a genuinely beneficial mutation drives selection while the rest of the sequence stays close to WT, making the variant hard to see against background without single-variant deconvolution. Either way, the high WT/WT-syn frequency understates the true adaptive diversity and argues for screening individual variants rather than trusting population-level sequencing. Consistent with this, re-integrating the passage 14 DAMGO-selected library into the fluorescence-screening strain and screening without sorting or 5-FOA counterselection raised the mutational load (NT 3.1, AA 2.2 vs. 1.6, 1.0) and revealed I300T to be 100% (Supplementary Fig. 8). This dominance likely reflects additional selective pressures, such as growth or expression, acting alongside signalling. It argues for some form of signal-focused screening after cell passaging at this early phase of biosensor evolution, though we reasoned that signalling-directed variants would come to dominate as evolution campaigns ran longer.

To test the signaling properties of the passage 14 variants directly, we transferred *OPRM1* variants from the single-copy clones onto high-copy 2μ plasmids and transformed them into a standard yeast biosensor background^15^, always against a wt-OPRM1 control. We chose DAMGO-selected variants as they had undergone more generations and compared arbitrarily selected early mutants to endpoint mutants. The results spanned a range of EC_50_ shifts relative to WT (1.44 μM): V286A/I300T/C348R and C13Y/I148T/D274G/I300T (passage 6) were both less potent (9.65 and 11.70 μM), while Y19H/S35P/R373T (passage 14) signaled modestly (4.02 μM) (Fig. 2E). Assayed against loperamide, the same three followed similar trends relative to WT (1.30 μM): 1.21, 4.60, and 2.76 μM, respectively (Fig. 2F). A second panel behaved differently: S30P/T69A/I300T (passage 6) and T155A/I300T/D383G (passage 14) gave DAMGO EC_50_ values of 2.68 and 4.74 μM against a WT of 2.13 μM, while V191A/D383G (passage 14) was roughly 12-fold more sensitive (0.17 μM) (Fig. 2G). The same rank order held under loperamide, with EC_50_ values of 0.71, 0.52, and 0.17 μM against a WT of 0.24 μM, although the DAMGO-selected V191A/D383G showed no appreciable improvement over WT (Fig. 2H). Compounding the fact that D383G was enriched after an additional ∼45 generations (while I300T showed a decrease), we hypothesize that D383G may be an example of ligand-specific signalling enrichment.

Taken together, OrthoRep-driven mutagenesis of OPRM1 yields distinct, host-tolerated signaling profiles. The ligand-specific enrichment we observe is consistent with the two agonists drawing out different mutational routes across independent populations, diversity of the kind useful for tuning biosensor dynamic range. Improved sensitivity, by contrast, was rare at this low-stringency selection regime which favours exploration over convergence. Reaching stronger, ligand-specific gains means sustaining selection over more generations, and this is where coupling growth to a distal reporter meets an intrinsic limit: over longer campaigns we saw a loss of divergence between the DAMGO and control groups, consistent with the emergence of suppressors (Fig. 2A). Sustaining evolution past escape is the purpose of the refresh strategy we introduce next.

### Selection escape motivates the use of a p1 landing pad to refresh evolution strains, establishing the HERO platform

Most reported OrthoRep applications place selection directly on the mutated protein, a single enzyme step, or mutational drift followed by enrichment screens^40-42^. Coupling OrthoRep instead to a distal reporter several steps downstream opens new failure modes. The clearest is selection escape (“cheaters”): during low-stringency passages (−His only), growth in the absence of both histidine and DAMGO took over after four passages (Supplementary Fig. 9). The same escape appeared in a static, non-mutagenized biosensor passaged alongside, which places the fault in the biosensor chassis rather than in OrthoRep-driven mutagenesis of the receptor (i.e. constitutive activation). Independent microscopy images of three escaper populations revealed elongated, shmoo-like cells which is a shape typical of *GPA1* disruption^19^. We found *GPA1-GNAI3* mutations in two of three isolated escaper clones, which may prevent Gαi3 from sequestering free G_β_/G_γ_ (Ste4 and Ste18) leading to sustained, aberrant signalling (Supplementary Fig. 10).

One approach we took to mediate this was adding an extra genomic copy of *GPA1-GNAI3* at a safe-harbor site, but this flattened agonist-induced signaling across the range we tested (Supplementary Fig. 11); we suspect the added Gαi3 shifts the Gα:Gβγ balance so that free Gβγ is re-captured by excess GDP-bound Gα after activation, dropping MAPK output below the reporter’s threshold, which is something seen in other studies^33,43^. Alternative strategies to eliminate host suppressors downstream of the entire cascade, such as 5-fluoroorotic acid supplementation in a biosensor driving a conditional *URA3* reporter, may also work in theory^44^, however this is more difficult to tune with the emergence of escape due to the toxicity of 5-fluorouracil paired with potentially leaky reporter expression^44^. Regardless, over many passages, other suppressor mutations outside this scope would inevitably appear.

Rather than querying each possible host suppressor mechanism, we adopt a simpler solution in this second phase of OrthoRep biosensor design. By adapting a system from Pisera *et al*., we address the challenge of moving coding sequences onto p1 copies of engineered yeast chassis, and use this approach to transfer the previously-mutated OPRM1 variants into a completely fresh biosensor background, thereby continuing rounds of evolution (Fig. 3A)^25^. We protoplast fused a p1 plasmid carrying a *URA3* marker flanked by attB sites into the 611-DNAP1 recipient, then supplied a curable plasmid encoding the TP901 recombinase. Previously-evolved *OPRM1*-*LEU2* cassettes, PCR-amplified with flanking attP sites, then recombine directly onto the p1 landing pad, exchanging *URA3* for the receptor and restoring leucine prototrophy. This refreshable, p1-based landing-pad GPCR biosensor, together with different mutational polymerases, is the platform we call HERO: in a single transformation step, continuous, user-defined mutagenesis begins or resumes under ligand-dependent growth selection.

**Figure 3.**
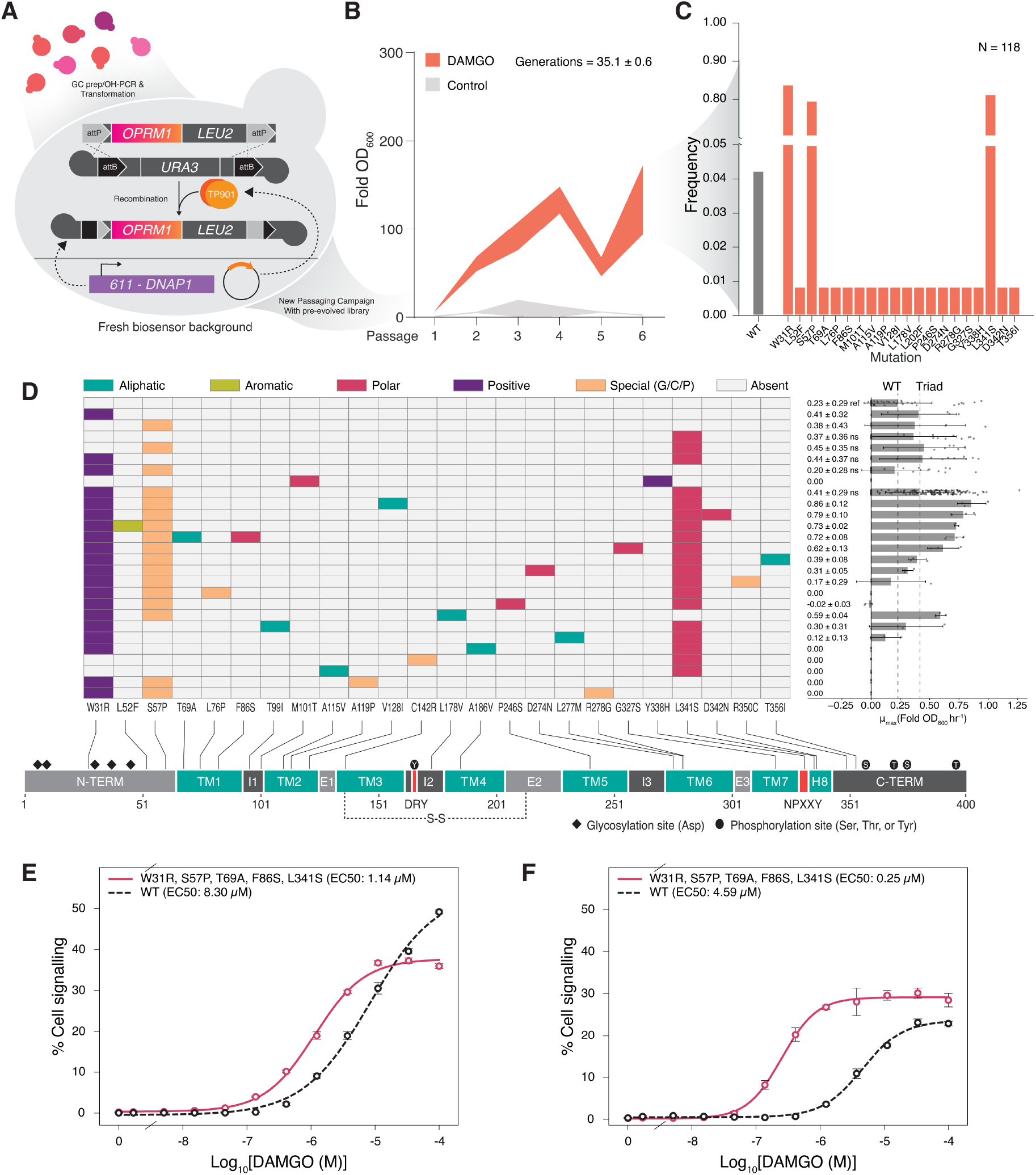
Selection escape and a modular p1 landing pad establish the HERO platform. **A** TP901-medited landing-pad refresh: a pre-evolved *OPRM1*-*LEU2* cassette with flanking attP sites recombines onto a *URA3*-marked attB landing pad in a fresh 611-DNAP1 biosensor background, restoring leucine prototrophy. **B** Growth (OD_f_/OD_i_) across six DAMGO passages after refresh, with estimated generations, agonist versus control. **C** Substitution frequencies among clones picked at random after 14 + 6 passages (N = 118); W31R, S57P, and L341S each reach roughly 0.8. **D** Per-position substitution map across the OPRM1 topology (N-terminus, TM1 to TM7, and helix 8, with the DRY and NPxxY motifs and glycosylation and phosphorylation sites marked), colored by amino-acid property; at right, per-clone 10 μM DAMGO-dependent growth rate (μ_max_, fold OD_600_ per hour, 48 h) for wild-type versus mutant clone at each position. **E** DAMGO dose-response of a representative winner (W31R/S57P/T69A/F86S/L341S) versus wild type on the ergosterol background and **F** on a cholesterol-producing background using a Log_3_ gradient, points are mean ± s.d.

We transferred the DAMGO-selected OPRM1 library from phase 1 (Fig. 2A) onto the p1 landing pad, selecting on SC-Leu. The variant library donor transformed into the 611-DNAP1 HERO strain at roughly 5 × 103 CFU/μg (Supplementary Fig. 12). After one round of SC-Leu + 5-FOA counterselection, we resumed passaging in 50 μM DAMGO in biosensor media (±3-AT) for six further rounds, consistent with our previous checkpoints (Fig. 3B, Supplementary Fig. 13). To check that library diversity carried over to the refreshed strain, we bulk-sequenced the pooled OPRM1 amplicons after recovery in non-selective media, and one DAMGO passage, comparing amino-acid substitutions against the full passage-14 set. Of 77 unique substitutions in the concatenated, clonal passage-14 data, 52 (68%) reappeared after refreshing; the pool also carried 591 additional substitutions in the assembled contigs (Supplementary Fig. 12), qualitatively suggesting a larger standing pool of diversity in our system than clonal sequencing at this scale resolves. Recovering 68% of prior substitutions indicates the refresh transfers most detectable diversity, with the 25 undetected substitutions being either below the detection limit of bulk pooled sequencing or lost in the transfer, the latter consistent with the lower CFU yield and possible selection during recovery.

We then picked and sequenced clones at random after integrating them as single copies into the growth-sensor strain. Three substitutions, W31R, S57P, and L341S, each appeared on their own in at least 80% of clones, 63% of which carried all three (Fig. 3C; Supplementary Fig. 14) and the WT/WT-syn share of the pool fell to about 4%. None of the three mutations had been seen in the pre-refresh clonal sequencing (passage 6 or 14); they surfaced only after refreshing and continued passaging, which we interpreted as a strong fitness benefit driving rapid enrichment. To set a screening threshold, we ran a DAMGO dose-response of WT-OPRM1 in the single-copy growth sensor in parallel and chose 10 μM DAMGO as a sufficiently high concentration to allow for detectable signalling, while being low enough to show improvements over WT (Fig. 1C; drug response of single-copy control strain). We then subjected every clone from the refreshed campaign to this concentration in triplicate. Clones carrying any or all of the triad grew roughly 2-fold faster (μmax) than the WT OPRM1 biosensor (Fig. 3D, right). No appreciable growth in the control group was observed in 24 hours.

Our screen recovered expected loss-of-function changes, a proline substitution at alanine 119 in TM2 (A119P) and loss of a conserved disulfide cysteine at position 142 (C142R), both of which cut drug-dependent growth (Fig. 3D). We expect these loss-of-function variants to reflect a standing pool of low-fitness mutants sustained by heteroplasmic winners, cross-feeding within the culture, or mutations arising shortly before library extraction. Separately, to test whether faster −His growth reflected receptor signalling rather than a generic growth benefit, we compared selective growth (10 μM DAMGO) with non-selective growth in synthetic complete medium for each variant, assayed in the fixed single-copy screening strain. The two were uncorrelated (Spearman ρ = 0.04, p = 0.66, n = 114; Supplementary Fig. 15). Because the variants sit in a common host background, this decoupling indicates that selective growth was not a readout of basal fitness; variants that grew well under selection were not simply those that grow well in general, supporting receptor signalling and expression rather than fitness basis for the enrichment.

To profile a representative improved variant in detail, we took a clone carrying the triad together with T69A and F86S (W31R/S57P/T69A/F86S/L341S) and measured its dose-response across 12 concentrations of DAMGO using plasmid-based expression. We note here that fold-improvement in sensitivity is the more stable metric across assays, since absolute EC_50_ values shift between experiments while the WT-normalized gain holds. In addition to signalling improvements, we also wanted to assess whether the substitutions were selected due to improved signalling specifically in the ergosterol membrane environments. On the ergosterol (native) background the mutant was 7.3-fold more sensitive than WT (EC_50_ 1.14 μM vs. 8.30 μM; Fig. 3E). The same plasmid expressed in a cholesterol-producing background, which partially mimics the mammalian membrane, was 18.4-fold more sensitive (EC_50_ 0.25 μM vs. 4.59 μM; Fig. 3F). Corresponding ENVY-induction histograms of both experiments showed a clearer delineation between on and off populations, where the wt-OPRM1 expressing plasmid had a leakier activation at the highest concentration of DAMGO, consistent with an improvement in the activation threshold of the otherwise weak DAMGO-OPRM1 biosensor interaction (Supplementary Fig. 16)^45^; on these bases the triad substitutions sharpen receptor signalling, and do not appear to be enriched as a condition of membrane composition^15^.

### The HERO platform generalizes across polymerases and receptors

Having established HERO in the second phase, we next expanded it. We first addressed screening. Plasmid-based expression carries three liabilities: copy-number variability, plasmid loss off selection, and heteroplasmy in pooled libraries^46^. We therefore adapted our stable, single-copy *pFIG1*-*HIS3* growth biosensor chassis to also possess a *pAGA1-ENVY* for tandem fluorescence reporting (Fig. 4A). With this improved schema, PCR-amplified libraries are integrated at single copy into the tandem reporter strain, and single clones can be sequenced, screened, and characterized simultaneously, removing multiple steps from our previous workflow. Integrated wt-OPRM1 DAMGO response curves for *HIS3*-dependent growth (24 h) and ENVY fluorescence (8 h induction) rose in proportion (Fig. 4B,C), confirming the two readouts agree. We used this dual-reporter strain exclusively from here forward.

**Figure 4.**
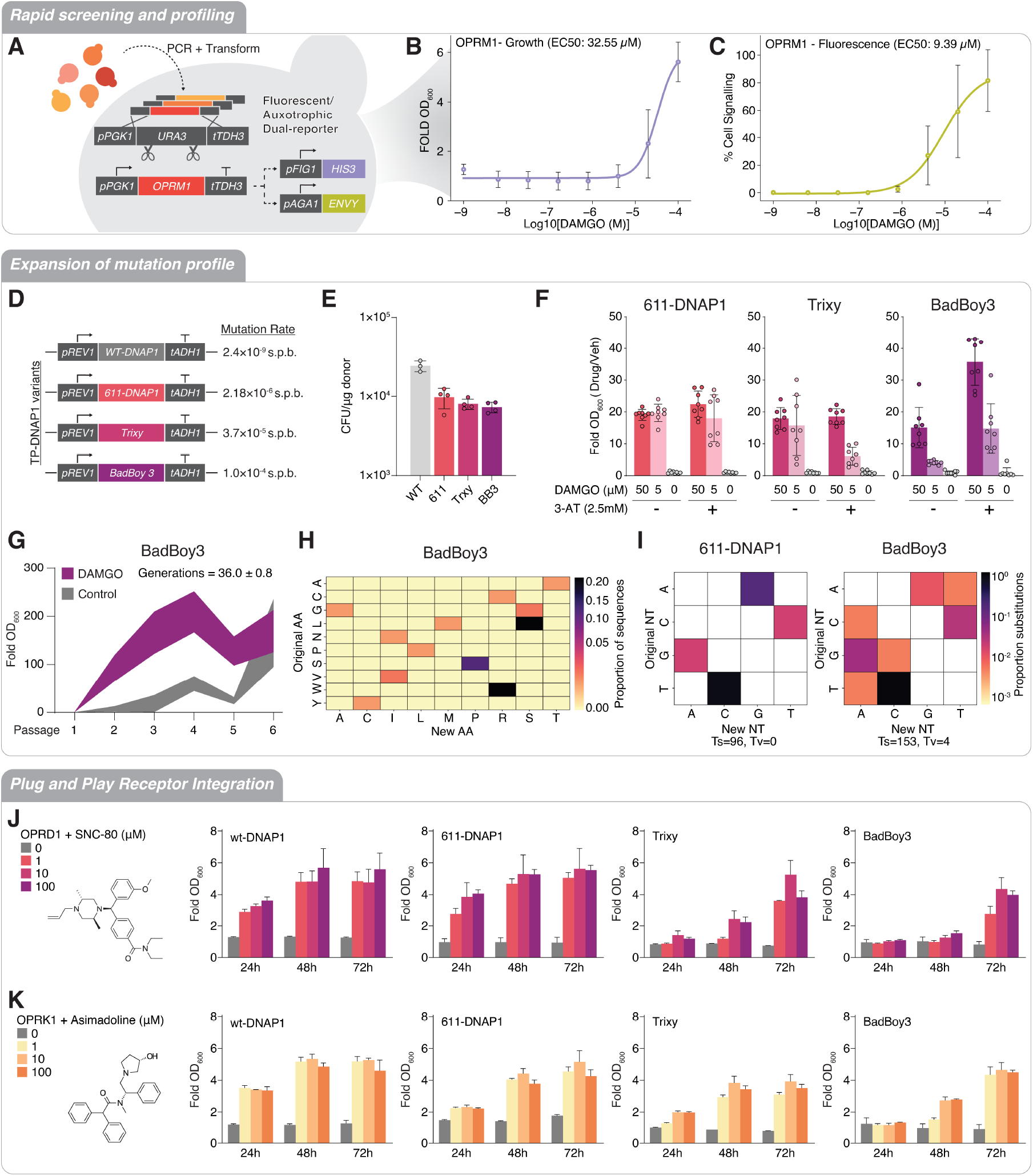
The HERO platform generalizes across polymerases and receptors. **A** The dual-reporter design: a pPGK1-URA3-tTDH3 landing pad receives a single receptor copy, read out by *pFIG1*-*HIS3* growth and *pAGA1-ENVY* fluorescence. **B** DAMGO growth dose-response and **C** DAMGO fluorescence dose-response of OPRM1-dual reporter (EC_50_ 32.55 and 9.39 μM), using a Log_5_ gradient, confirming that the two readouts agree. **D** TP-DNAP1 variants and their per-base mutation rates (wt-DNAP1 2.4 × 10^-9^, 611-DNAP1 2.18 × 10^-6^, Trixy 3.7 × 10^-5^, BadBoy3 1.0 × 10^-4^ s.p.b.). **E** Viable CFU across the four polymerases. **F** DAMGO growth with and without 2.5 mM 3-AT for 611-DNAP1, Trixy, and BadBoy3. **G** Growth across six DAMGO passages for BadBoy3, agonist versus control. **H** BadBoy3 amino-acid substitution spectrum. **I** Nucleotide substitution spectra for 611-DNAP1 (Ts = 96, Tv = 0) and BadBoy3 (Ts = 153, Tv = 4), showing the transversion bias of BadBoy3. **J** Plug-and-play expression of OPRD1 with SNC-80 and **K** OPRK1 with asimadoline, measured as fold OD_600_ at 24, 48, and 72 h across all four polymerases. Bars are mean ± s.d.

Next, we reconcile the limitations of 611-DNAP1 as the sole mutagenic driver as it possesses lower processivity and bias towards nucleotide transitions that might be required^5^. To expand the evolution profiles and speeds we added two polymerases from Rix *et al*. to the landing-pad biosensor platform from Fig. 3A^5^. These were chosen primarily for their increased mutation rates of 3.7 × 10^-5^ (Trixy) and 1.0 × 10^-4^ s.p.b. (BadBoy3), against 2.4 × 10^-9^ (wt-DNAP1) and 2.18 × 10^-6^ (611-DNAP1) (Fig. 4D), placing BadBoy3 roughly 46-fold above 611-DNAP1, and because both canonically show a stronger transversion bias that should expand the mutational landscape reached. Upon integration onto the p1 landing pad, the higher error rate tracked loosely with lower viable CFU when comparing to the wt-DNAP1, however no significant difference existed among the three mutagenic polymerases (Fig. 4E). Additionally, all three mutagenic strains gave the expected DAMGO growth responses with and without 2.5 mM 3-AT (Fig. 4F). We then ran a 48-h cell passaging campaign under the same conditions used for the 611-DNAP1 refresh campaign (Fig. 4G; Supplementary Fig. 17). After only six rounds, the BadBoy3/DAMGO pool had converged on the same W31R/S57P/L341S positions recovered from the 14 + 6-round refreshed 611-DNAP1 campaign (W31R 24%, L341S 21%, S57P 12% of clones; Fig. 4H).

Because polymerase replication rate governs variant supply but not the selection that drives fixation, the ∼46-fold faster BadBoy3 generates these variants sooner, reaching the same convergent targets in less than a third of the generations (∼36 generations over six passages vs. ∼123 over the 14 pre-refresh plus six post-refresh passages using 611-DNAP1); consistent with this, the population sits mid-sweep, with the triad enriching (8% of clones already carrying all three) but not yet dramatically enriched. Although the mean mutation frequency was virtually identical to 611-DNAP1 (1.2 vs. 1.3 NT substitutions per clone; Supplementary Fig. 6 and 18), and the triad had not yet enriched, we did observe an improvement of transversion frequency. Compared to 611-DNAP1 over roughly the same number of generations, BadBoy3 introduced transversions where 611-DNAP1 produced essentially none (Tv = 4 vs 0, Ts = 153 vs 96; Fig. 4H,I). 611-DNAP1 is strongly transition-biased, so it cannot reach substitutions that require a purine-pyrimidine change, regardless of generation count. Three BadBoy3 substitutions are of this type: N193I, G269A, and L392M. Each requires a transversion at the codon level and has no transition-only path, and none appear in the 611-DNAP1 variant pools. These variants were recovered by sequencing but not phenotype, so they mark newly reachable sequence space rather than confirmed functional gains. G269A is notable in that it arose alongside L341S, placing a transversion-only change adjacent to the convergent triad. The choice of polymerase therefore controls which regions of sequence space an evolution campaign can reach.

Lastly, we show that HERO extends to other opioid receptors without redesign. Using a *LEU2*-marked PCR donor for OPRD1 (Fig. 4J) and OPRK1 (Fig. 4K), we observed functional expression over 24-, 48-, and 72-h time courses across all four polymerases. Signal for Trixy and BadBoy3 was lower than for the wt- and 611-DNAP1 OPRM1 strains, which may reflect a larger fraction of mutated p1::GOI copies or lower processivity and copy number. A six passage OPRD1 campaign under 611-DNAP1 against OPRD1-specific agonist, SNC-80, (Supplementary Fig. 19) recovered a recurrent Ala-to-Thr substitution (A51T) alongside a small number of additional changes. These were broadly consistent with the polymerase’s transition bias, tended to be conservative swaps between chemically similar residues, and appeared distributed across the receptor rather than clustered at the presumed ligand pocket. This pattern of single AA mutations being favored early on is consistent with our previous OPRM1 campaigns, where selection may be a result of expression, folding, and/or agonist contacts. We re-evaluated the utility of screening individual variants of this campaign, as OPRD1 is already highly sensitive to the benchmarked SNC-80, and this pairing offered little room to improve. We instead looked for a weaker receptor/ligand interaction using OPRD1 as a proof of concept for evolving improved signalling.

### High-throughput evolution of OPRD1 sharpens a weak agonist interaction

Continuous directed evolution for improved selected outcomes needs a starting interaction, however small, that the system can detect. To find an evolvable handle for OPRD1 OrthoRep evolution, we assayed the OPRD1-HERO strains against six commonly studied agonists in the HERO strains themselves (Fig. 5A,B). SNC-80 gave a strong response, but loperamide sat near the detection floor in the wt-DNAP1 strain, and lower still in the mutagenic 611-DNAP1 strain: this weak, but detectable loperamide response was a suitable interaction we sought to improve in a 24-well plate, automated evolution campaign.

**Figure 5.**
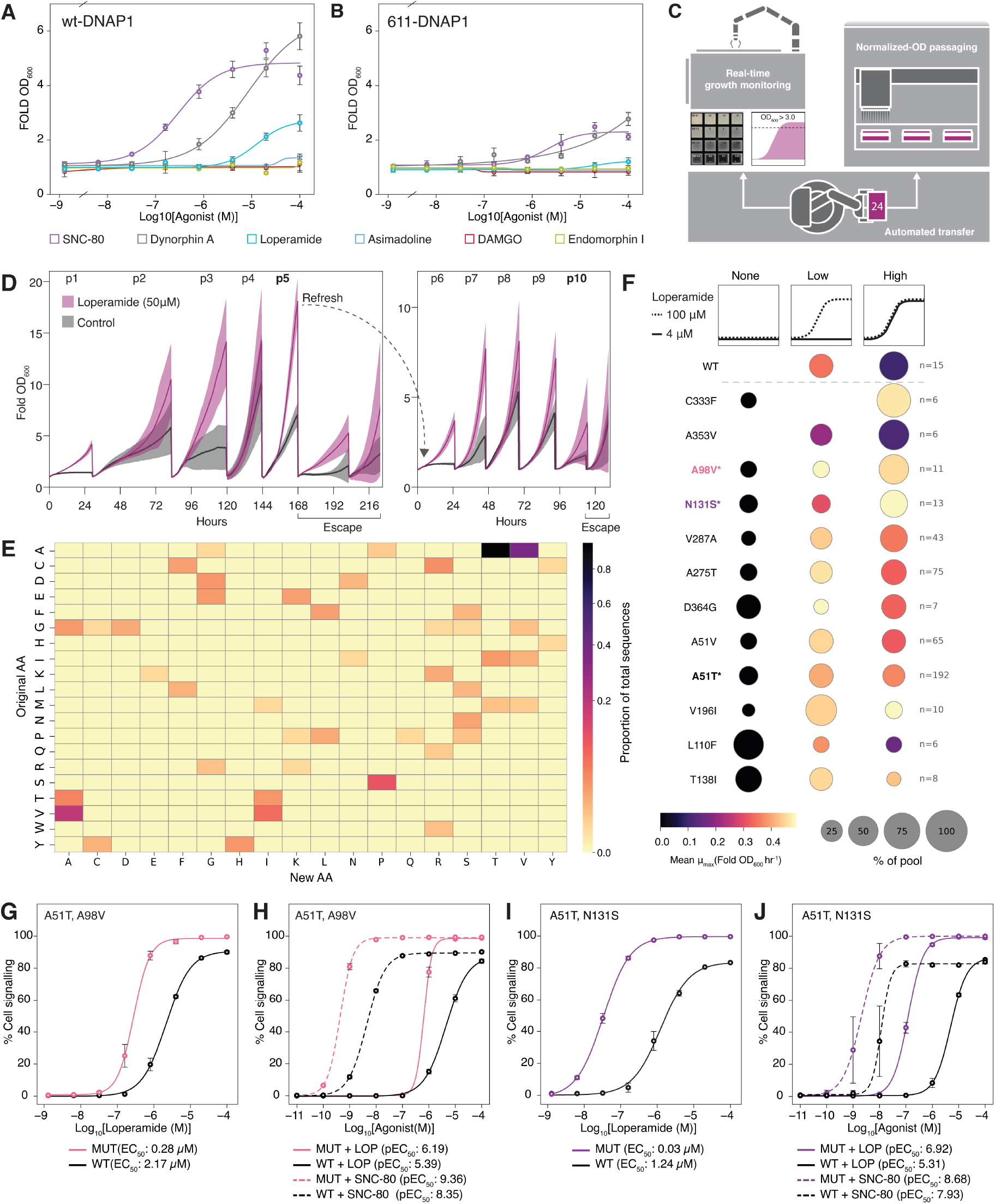
High-throughput evolution of OPRD1 sharpens a weak agonist interaction. **A** Growth dose-responses of the OPRD1-dual reporter strain in the wt-DNAP1 and **B** 611-DNAP1 backgrounds against six agonists (SNC-80, dynorphin A, loperamide, asimadoline, DAMGO, endomorphin I). **C** The automated normalized-OD passaging platform, which monitors growth in real time and transfers each of 24 cultures once it crosses OD_600_ > 3.0. **D** Automated loperamide (100 μM) versus control across ten passages, with a background refresh after passage 5. **E** Amino-acid substitution matrix after 10 passages **F** Sensitivity classes of the most frequent OPRD1 substitutions (rows; ≥6 carriers each), with WT reference (top). Each carrier genotype is binned by loperamide response clearing a classification gate: none (no growth in any category), low (100 μM growth only), or high (4 μM and 100 μM growth). Dot size is fraction of carriers in that class; color is mean μmax of carrier within a sensitivity class. Asterisks indicate substitutions assayed in panels G to J. **G** Loperamide dose-response of A51T/A98V versus wild type using a Log_5_ gradient and **H** the same pair against loperamide and SNC-80 using a Log_10_ gradient and expressed in pEC_50_. **I** Loperamide dose-response of A51T/N131S versus wild type using a Log_5_ gradient and **J** the same pair assayed against loperamide and SNC-80 with a Log_10_ gradient and expressed in pEC_50_. Points are mean ± s.d.

In this third phase, we evolved OPRD1 against loperamide on an automated, normalized-OD passaging platform that monitors growth in real time and transfers cultures once one of the 24 populations crosses an OD_600_ > 3.0, at which point, all cultures are normalized to the same OD_600_ and passaged again (Fig. 5C). At this threshold, fast-growing cultures did not reach saturation while slower lineages had additional exponential growth time before the next passage (Fig. 5D). We reasoned that OD_600_ thresholds lower than 3.0 for the top-performers could bottleneck these slower lineages more severely at each passage, risking stochastic loss of variant populations we intended to track. The automated evolution system was comprised of a Growthprofiler 960 (Enzyscreen) for real-time growth monitoring and a Lynx liquid handler (Dynamic Devices), coupled with a spinnaker arm for plate transfer between passaging steps (Thermo). Over ten automated cell passages, loperamide-treated lineages pulled steadily away from the controls, with a background refresh after passage 5 and full escape emerging after passage 10 (Fig. 5D). Long-read consensus sequencing of individual clones showed the mutational load climbing from passage 5 to passage 10 (NT mean 3.7 to 4.6; AA mean 1.8 to 2.4; Supplementary Fig. 20). In addition to the aggregate rise in mutational frequency, well-level resolution across the 24 populations (Supplementary Fig. 21) revealed reproducible outcomes: A51T was present in all 24 passage-5 populations, and A51V and A275T recurred across independent lineages, evidence of parallel evolution, alongside substitutions unique to individual populations, such as N131S, V59I, and A353V (wells C3, C4, and D2).

Following genotype analysis, we ran a growth-based dose-response of the wt-OPRD1 dual reporter control to determine optimal single-dose for deconvolution of clone function; SNC-80 reached an EC_50_ of 0.09 μM against loperamide’s 18.19 μM (Supplementary Fig. 22). From this curve, we selected 4 μM of loperamide as a limiting, discriminating concentration and 100 μM as a permissive concentration at which wt-OPRD1 responds. Decomposing the evolved pool into individual substitutions (Fig. 5E) and classifying clones by loperamide sensitivity (Fig. 5F) separated the population into three response classes, none (neither concentration), low (100 μM only), and high (both 4 and 100 μM), revealing which substitutions carry the phenotype. A51T was by far the most abundant substitution (n = 192), followed by A275T (n = 75), A51V (n = 65), and V287A (n = 43); yet carriers of these common substitutions fell across all three response classes, whereas the rarer C333F, A353V, and A98V sat predominantly in the high-sensitivity (4 μM) class. Pool variant abundance therefore did not track sensitivity, marking the frequent substitutions as permissive background changes rather than primary drivers of the sensitized phenotype. A353V, though predominantly a high-class responder, carried a comparatively low average μ_max_ at 4 μM, and while C333F was one of the strongest hits, it gave only modest shifts when validated in two separate combination mutants (Supplementary Fig. 23). Because the discrete screen scores whether a clone clears the limiting-dose gate and how well it grows there rather than the magnitude of its EC_50_ shift, a top-ranked hit can still validate modestly.

Fluorescence-based dose-response curves revealed A51T/A98V and A51T/N131S as having the largest confirmed potency gains. A51T/A98V shifted the loperamide EC_50_ to 0.28 μM from a co-assayed WT of 2.17 μM (Fig. 5G). The same mutant also sharpened the response to SNC-80 across a 10-fold gradient, raising the pEC_50_ from a WT of 8.35 to 9.36 (Fig. 5H). A51T/N131S followed the same pattern more strongly: it dropped the loperamide EC_50_ to 0.03 μM from a WT of 1.24 μM, roughly a 40-fold gain (Fig. 5I), and raised the SNC-80 pEC_50_ in a 10-fold gradient from a WT of 7.93 to 8.68 (Fig. 5J). Additional isolated mutants showed the same trend of improved loperamide potency, and all displayed EC_50_s below the WT controls (Supplementary Fig. 23). These substitutions did not simply retune the receptor for loperamide. Both A51T/A98V and A51T/N131S mutants also gained SNC-80 sensitivity, broadening the receptor’s response across chemically distinct ligands rather than loperamide alone. The strongest SNC-80 shift, from A51T/A98V, reached the sub-nanomolar range of roughly 0.4 nM in the co-assayed run. However, because each run’s absolute EC_50_ depends on its co-assayed wild-type standard, we report these improvements as fold-shifts over the paired WT rather than as absolute affinities; mutant EC_50_ fold-shifts relative to WT were highly reproducible across independent 5-fold and 10-fold loperamide dilution gradients (loperamide gains: 7.8-vs 7.0-fold, Fig. 5G and H respectively; 41.3-vs 41.2-fold, Fig. 5I and J respectively), confirming that potency shift is a reliable measure^47^.

Together, these results show that HERO, combining a refreshable host, a fixed single-copy dual reporter, and a choice of polymerase, can take a modest GPCR-agonist interaction and, with minimal human intervention, improve its sensitivity by roughly 40-fold.

Notably, the same substitutions generalized across agonists rather than specializing: A51T mutants gained sub-nanomolar SNC-80 sensitivity while remaining strong loperamide responders, indicating general improvement in receptor function rather than a trade-off between ligands.

## DISCUSSION

We have succeeded in adapting OrthoRep’s continuous *in vivo* mutagenesis into a generalizable, first-generation system for engineering GPCRs. HERO overcomes the bottleneck of introducing GPCR sequences into the OrthoRep biosensor chassis, then passages them over many generations. Using the p1 landing pad to move mutant libraries into uncompromised backgrounds or different mutagenic chassis solves the recurring issue of escape instead of relying on protoplast fusion, homology-directed repair or extensive MAPK modifications, as shown in the phases of this project. Refresh need not be a one-time step; it can be repeated multiple times and is expected to broaden the explored sequence space each cycle, as seen between passages 5 and 10 of the OPRD1 campaign. Our screening system also facilitated population *and* individual-mutant characterization in tandem. All this, combined with the automated normalized-OD platform (Fig. 5C,D), is a step closer to continuous evolution of GPCRs becoming routine, demanding neither iterative library preparation nor extensive rational legwork^38^.

Directed evolution and protein engineering aim to evolve toward arbitrary functions, which ultimately hinges on selection. For enzymes this is frequently the limiting step: directed evolution is tractable when high-throughput screens or selection exist, but otherwise impractical. Bespoke GPCR biosensors tuned to specific metabolites offer such selection and become recursive inputs for continuous evolution of difficult-to-engineer enzymatic pathways expressed in the same biosensor host, or co-cultured cross-feeding strains. Evolvable pairs of sensors and key intermediates in heterologous hosts include opioid receptors and (S)-reticuline downstream products^18,48^, or cannabinoid receptors^49^ and the production of useful cannabigerolic acid derivatives^50,51^.

While metabolic engineering remains a primary aim, HERO has enabled basic inquiry into GPCRs in heterologous hosts and their structure–function relationships^52,53^. GPCRs, although ubiquitous, are more constrained in potential evolutionary directions due to the need to accommodate ligand interactions, membrane environments, extracellular and intracellular environments, and the downstream signalling partners^38,54^. W31R, S57P, and L341S recurred across independent campaigns, indicating positive selection rather than drift, yet none fall in the orthosteric pocket or at the G-protein interface: W31R and S57P lie in the N-terminal region and L341S at the TM7-H8 junction near the NPxxY motif (OPRM1, UniProt P35372)^55^. In the ergosterol background in which they were selected, this triad contributed to gains in sensitivity up to ∼7-fold over wild type (Fig. 3D-F); the same variants reached ∼18-fold in a cholesterol-humanized background they had never evolved against. Because these gains are not specific to membrane-composition nor the orthosteric pocket, and because heterologous GPCR evolution tends to enrich folding and stability mutations over ligand-recognition ones^9,38^, we favour effects on receptor stability or functional expression as the likeliest explanations, though these remain to be tested in our system. This trend appears in OPRD1: the A51T/A98V double mutant improved response to both loperamide and SNC-80 rather than trading one for the other (Fig. 5G–J), and neither residue maps to reported orthosteric contacts^56^.

The only mutation we characterized with prior links to signalling is N131S, which coordinates the conserved allosteric sodium pocket position. This substitution alters OPRD1 signalling and stabilizes the active-like state in mammalian models^57,58^. N131S was recovered only alongside A51T, the pair conferring the largest loperamide improvement, so we cannot yet isolate its individual contribution, and the basis of this asparagine-to-serine change remains unresolved in our yeast biosensor. That HERO surfaces functional variation in these distributed, structurally under-resolved regions is a strength; that it has not yet delivered pocket-level, ligand-specific gains points to the obvious next step.

Different selection schemes could help towards reaching such discrete functional substitutions^42,59^. Passaging without 3-AT prioritized mutational accumulation under less-stringent selection with the aim of accruing greater diversity and de-prioritizing functional escape^60,61^. The near saturating concentration of ligand carries two potential caveats: first our assay stringency is influenced by potential metabolic cross-feeding, since histidine produced by signaling cells might sustain their neighbors to an appreciable degree; deleting the histidine importers *HIP1* and *TAT1* may alleviate this^62,63^. Second, near-saturating ligand compresses the dynamic ceiling for improvement of signalling which may allow non-optimal alleles to persist across p1 copies, which is the expected cost of diversity that our low-stringency design intends. Passaging OPRM1 with 3-AT as a means of tuning this stringency produced differences in substitution count, but substitutions unique to 3-AT in this campaign did not appear enriched (Supplementary Fig. 8 and 19, Source data for Fig. 2). Interestingly, beginning the loperamide-OPRD1 campaign with an N-value of 24 separate populations increased representation of unique genotypes within individual wells, despite evolving under identical selection conditions^64^, and some of these unique mutants displayed putative functional gains (Fig. 5G-H, Supplementary Fig. 21 and 23). Other continuous-evolution systems have ratcheted stringency by tapering a fed metabolite until the evolving enzyme supplies it^42^; dynamically lowering ligand concentration and running multiple evolution concentrations in parallel^59^ is the subject of future work.

Complementary to selection regimes, we discuss screening methods. Our initial FACS sorting scheme showed depleted I300T representation and increased the relative frequency of D383G, but it also increased the share of wt-OPRM1. This is consistent with other GPCR biosensor work detailing how sorting can impose selection pressure different from the intended output, however this can be mitigated with model-guided approaches to determine optimal gating^65^. Growth-coupled readouts reliably report signalling strength and recover functional variants in yeast GPCR systems^66^. In our hands, random clone-picking followed by high-throughput growth assays proved informative, as it allowed mapping of permissive as well as deleterious positions. With regards to single-variant characterization of mutants, plasmid expression is not ideal as it introduces reservoirs of off-state cells and, when performing pooled transformations, heteroplasmy confounds the link between genotype and phenotype^67^. We therefore adapted the single-copy, growth-based chassis with a tandem fluorescent reporter, enabling population- and single-variant-level drug-response characterization from the same transformation^15^. With HERO now established, modifying stringency, screening, and the biosensor chassis itself are incremental tasks we intend on pursuing.

A final consideration for improving sensitivity and function is that, unlike one-shot library methods which begin already diversified^68,69^, continuous *in vivo* evolution in our hands started from a WT sequence. About half of sequenced clones stayed WT or synonymous-WT after ∼90 generations of selection (Supplementary Fig. 6 and 7). These persistent WT and low-fitness pools are explained by the variable copy number and heteroplasmy of p1::GOI constructs (Fig. 3D): under multicopy heteroplasmy, functional copies may mask the fitness cost of WT, synonymous, and non-functional alleles sharing the cell, allowing them to persist^39^. Additionally, the mutational profiles afforded by DNAP1s further limit what we can see: because transitions dominate the 611-DNAP1 spectrum, the recovered substitutions cluster into a smaller set of accessible residue swaps, as seen with recurrent OPRD1 A to V/V to A symmetry in both SNC-80 and loperamide campaigns. Ultimately, this bias warranted our characterization of BadBoy3, however equalizing substitution bias does not eliminate the hurdles intrinsic to starting evolution in a defined codon neighborhood. Rather than being a competitor, HERO can complement other directed evolution techniques: integrating scanning site-saturated^70,71^, computationally designed^72,73^, or EP-PCR amplified^38^ GPCR libraries onto the p1 landing pad of the HERO biosensor provides access to codon space and combinatorics otherwise underexplored by our current system. Deploying HERO passaging thereafter would allow users to resolve the combinatoric gaps necessary to further satisfy the defined selection constraints.

## ACKNOWLEDGEMENTS

We thank the Concordia Genome Foundry members Nick Gold, Angela Quach, and Mindy Melgar for assistance with automation and high-throughput screening, and Manpreet Vilkhu for providing MVY09 and sharing insights from her single-copy screening work. We thank Alireza Tanoori, Jacob Hambalek, Rory Williams, and Olek Pisera for extra guidance in designing HERO and providing strains at the University of California, Irvine. This study was financially supported by an FQRNT Team grant, and an NSERC Discovery grant. C.B.D. was supported by an NSERC Graduate Student Fellowship, a MITACS Globalink fellowship, and a Lallemand Fellowship, B.D.M.B. was supported by a Concordia University Horizon Postdoctoral Fellowship. V.J.J.M. is supported by a Concordia University Research Chair. M.W. is supported by a Canada Research Chair. C.C.L. is supported by the Institute for Rapid Antibody Engineering and Evolution as part of the Engineering+ Health Initiative of the UCI Samueli School of Engineering.

## AUTHOR CONTRIBUTIONS

C.B.D., B.D.M.B., C.C.L., M.W. and V.J.J.M. designed the research.

C.B.D. performed the experiments, and D.M. assisted with delta- OR/SNC-80 evolution campaigns and sequencing. O.R. and B.D.M.B. constructed donor and recipient chassis strains for phase 1 OrthoRep biosensors. F.A. and C.B.D. established the automated passaging workflow. V.J.J.M. and M.W., and C.C.L. supervised the research.

C.B.D. wrote the manuscript, with editing help from B.D.M.B., C.C.L., M.W., and V.J.J.M.

## DATA AVAILABILITY

All source data, including raw values, are provided in the Source Data file. Raw sequencing data can be found under the NCBI accession PRJNA1533496.

## CODE AVAILABILITY

Code used for drug response curves, demultiplexing, mutational analysis, and automated passaging can be found at https://doi.org/10.5281/zenodo.22903774.

## ADDITIONAL INFORMATION

Supplementary information is available in the online version of the paper. Correspondence and requests for materials should be addressed to V.J.J.M.

## METHODS

### General strain and plasmid construction

Yeast strains and plasmids used in this study are summarized in Supplementary Tables 1 and 2, respectively. Biosensor strain chassis and opioid receptor sequences are derived from Bean *et al*^*15*^, and the p1 landing pad donor was derived from Pisera *et al*.^25^. For all genomic integrations, pCas9 plasmids from the Mylo-Toolkit and Dykstra *et al*. were used with targeting hybridized oligonucleotides^74,75^. Amplicons for constructs and chassis integrations were generated using Phusion HF polymerase (Thermo), with the exception of library refreshing. Plasmids were assembled via Golden Gate, homology directed repair in yeast, or splice overlap extension (SOEing) and transformed into DH5α *E. coli*^75-77^. Primers were synthesized by Thermo, except for otherwise noted IDT primers. All relevant primer pairs are listed in Supplementary Table 3. Clonal plasmids were sequence verified by whole-plasmid sequencing (Plasmidsaurus).

### Chemical transformation and background refreshing

Yeast transformations were carried out using an adapted version of the Gietz method; Overnight yeast culture grown to an OD600 of 1.2, was pelleted and washed with distilled water followed by 100 mM LiOAc solution. Pellets were then resuspended in 150 μL transformation solution (PEG, 33.3%; lithium acetate, 100 mM; denatured salmon sperm DNA 300 ng μL-1) containing 200 - 500 ng of pCas9 plasmid and/or 500 – 2000 ng of linear donor DNA. Samples were heat-shocked at 42°C for thirty minutes prior to 24-hour recovery in 600 μL of YPD broth. Transformants were selected using media containing geneticin (G418, 200 ng μL-1) and cured on YPD media. Background refresh was conducted using the same transformation technique, with the exception of using KOD One Blue Master Mix (Toyobo) PCR product without purification directly as a donor into the HERO LP strain.

### OrthoRep biosensor donor and recipient construction

Receptor–marker transcriptional units were assembled by Golden Gate cloning of each opioid receptor coding sequence into an integrating p1 vector carrying the 10B2 p1 promoter, a GFP drop-out cassette, an A75 repeat region, an Rz self-cleaving element, and a *LEU2* marker^42,76,78^. For the initial OPRM1 donor strains, the integrating vector was linearized and transformed into the p1/p2-bearing strain GA-Y319, and p1::OPRM1–*LEU2* integrants were selected on synthetic complete medium lacking leucine (SC–Leu; Supplementary Fig. 1B,C). The final HERO donor strain was generated by disrupting *TRP5* in a strain carrying the p1::*attB*–*URA3*–*attB* landing pad, yielding a *trp5*Δ0 auxotroph. Recipient strains were derived from BBY1640^15^ biosensor and expressed OrthoRep DNAP1 variants from a *pREV1*/*tADH1* promoter–terminator pair integrated at the *FF18* locus^79^.

### Protoplast fusion

Donor and recipient strains were combined by protoplast fusion^2,4,59^. Strains were grown to saturation overnight at 30 °C in the appropriate selective medium; where additional biomass was required, cultures were grown overnight in 3 mL, back-diluted 1:33–1:50 into 5 mL, and grown to saturation a second time. Cells were pelleted (4,000 × *g*, 2 min), resuspended in 500 μL dH_2_O, transferred to microcentrifuge tubes, and re-pelleted (3,000 × *g*, 1 min). Equal wet-pellet masses of donor and recipient (∼12.5 mg each, corresponding to ∼3.75 OD_600_ mL of the p1::GOI donor or ∼5 OD_600_·mL of the BY4741-derived recipient) were used per fusion. Each pellet was resuspended in 75 μL of 0.06 M EDTA / 0.2 % (v/v) β-mercaptoethanol, donor and recipient suspensions were combined (150 μL final), and the mixture was incubated at 30 °C for 30 min, inverting to mix at 15 min. Cells were resuspended gently and then pelleted (400 × *g*, 1 min; used for all subsequent spins unless noted), washed once in 250 μL 0.6 M KCl, and resuspended in 400 μL Buffer I (citrate–phosphate buffer pH 6.1, 0.6 M KCl, 0.01 M EDTA) containing zymolyase at 6 U/mL. Digestion proceeded for 1 h at 30 °C with periodic agitation. Protoplasts were then pelleted and resuspended in 400 μL freshly prepared Buffer II (33 % (w/v) PEG-3350, 0.6 M KCl, 50 mM CaCl_2_) and incubated at 30 °C for 30 min with occasional agitation. Fused cells were pelleted (700 × *g*, 2 min), resuspended in 400 μL Buffer III (0.6 M KCl, 50 mM CaCl_2_), and 200 μL (range 20–400 μL) was mixed by inversion into 15 mL of molten regeneration medium (SC -Leu/Trp agar containing 3 % (w/v) bacto-agar and 0.6 M KCl, held at ∼45 °C) in a 15 mL tube. The suspension was poured immediately and fusants were selected at 30 °C after 5 days.

### Propidium iodide staining

Ploidy was assessed by propidium iodide staining and flow cytometry^80^. Cells were grown overnight at 30°C in 1 ml YPD in deep-well plates alongside control strains of known ploidy, then diluted to OD600 ≈ 0.6, pelleted (500 × g, 3 min), and fixed in 1 ml 70% ethanol for at least 1 h. Fixed cells were washed twice in 50 mM sodium citrate, sonicated briefly, and treated with 0.5 mg/ml RNase A in 50 mM sodium citrate/40% glycerol at 37°C for 2 h. Cells were stained with propidium iodide (25 μg/ml final) in 50 mM sodium citrate overnight at 37°C in the dark, keeping the PI concentration constant across all samples and experiments to allow comparison. Immediately before acquisition, samples were sonicated and diluted 1:40 into staining buffer (25 μg/ml PI, 50 mM sodium citrate), and 20,000 events were collected per sample. Unstained cells served as a negative control. This was performed for the wt-, 611-, and 1-4-2-DNAP1 strains.

### OrthoRep biosensor passaging

For manual OPRM1 passaging campaigns, biosensor strains were grown to saturation in non-selective media for 16 hours, washed in dH2O, and inoculated to a starting OD_600_ of 0.3 in biosensor media (synthetic dropout media without histidine or leucine, 100 mM TRIS-HCL, pH7.1). Cultures were split into 2mL replicates (n=4) and supplemented with either agonist (DAMGO or loperamide, 50 μM) or dimethyl sulfoxide at the same volume as a passage step control. After 48 h of growth at 30 °C with shaking at 300 rpm, OD_600_ was recorded and the agonist-treated replicates were back diluted 1:100 for OPRM1, or 1:50 for OPRD1, into fresh biosensor media with/without agonist and grown for a further 48 h. Each passage was archived in 25 % glycerol and frozen, preserving timepoints in the event of escape or for later mutational analysis. OPRD1 manual passages with SNC-80 were performed similarly, except that SNC-80 was used at concentration of 5 μM and a wash step was introduced at each passage to limit drug carry-over into the control groups. For each passage, the OD_600_ at inoculation was taken as the previous passage’s final OD_600_ divided by the dilution factor. The fold change in OD600 and the number of generations per passage were then calculated as:

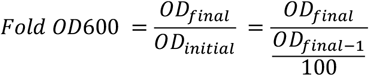

Using this, generations were estimated by the following calculation:

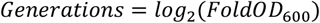

### Stable mutant library screening

Mutant GPCR libraries were integrated into screening strains carrying a *pPGK1–URA3–tTDH1* landing pad at the *FF16* locus^79^, together with either a *pFIG1-ENVY* fluorescent reporter, a *pFIG1-HIS3* growth reporter, or a *pFIG1-HIS3* /*pAGA1-ENVY* dual reporter; these strains were used for clonal sequencing and phenotypic assays. Purified, or unpurified KOD One Blue Master Mix library donor amplicons carrying 50-bp homology arms were co-transformed with a Cas9 plasmid (pCas9) expressing a dual gRNA targeting *URA3*, driving replacement of the landing pad with the integrating mutant receptor. Following transformation, cells were either recovered for 24 h in YPD and plated on G418 + 5-FOA or passaged once in liquid G418 + 5-FOA after the same recovery period; loss of *URA3* (5-FOA resistance) reported on successful gene disruption at the landing pad locus. For Figure 2, the top 30 % of the wild-type-ON population was collected by FACS after 8 hours of 50uM agonist induction, plated, and individual colonies were picked for sequencing. For all other evolution campaigns, colonies were picked at random into 96-well plates using a QPix colony picker (Molecular Devices) and propagated for phenotypic testing and clonal sequencing. Figures 4 and 5 used an improved tandem-reporter biosensor that combined the growth and fluorescent readouts, allowing growth-based screening and fluorescence-based drug-response curves to be performed in a single step.

### Sequencing mutant libraries

Sorted or randomly picked colonies were boiled for 10 min in 50 μL of 40 mM NaOH and diluted 1:4. Barcoding primers annealing either at the immediate start and end of the receptor CDS or ∼30 bp up- and downstream were designed using 7-mer barcodes from Rix *et al*.^5^ (Supplementary Table 3). Primers were dispensed with an Echo acoustic liquid handler to a final concentration of 0.5 μM into 20-μL reactions of KOD One Blue Master Mix containing 5 μL of boiled template. Cycling conditions were: 98 °C, 10 sec; 57 °C, 5 sec; 68 °C, 25 sec for 32 cycles, with a final extension of 5 minutes at 68 °C. Ten μL of each barcoded reaction was pooled and submitted to Plasmidsaurus or FLOW Genomics for PCR amplicon sequencing. Every sequencing run included a spiked, barcoded, unmutated wild-type receptor sequence as a control.

### Mutational analysis

Demultiplexing, consensus generation, and mutation calling were performed with a custom pipeline inspired by Maple from Rix *et al*.^5^. Reads were returned from Plasmidsaurus which uses R10.4.1 flowcells (Oxford Nanopore Technologies). They were then demultiplexed in two passes with Dorado [ver. 0.9.6]^81^: first by the forward 7-mer barcode, then, after reverse-complementation, by the reverse barcode, using custom barcode arrangements and --no-trim. Reads shorter than 1,200 bp were discarded prior to the reverse pass, and forward/reverse barcode pairs were resolved to well identities; wells with fewer than 10 reads were excluded. Reads for each well were aligned to the receptor reference with minimap2^82^, sorted and indexed with samtools^83^, and a per-well consensus was generated with Medaka (version 2.2.0)^84^. The coding sequence was extracted from each consensus by anchoring on the receptor start codon and trimming in-frame to the reference length, then globally aligned to the reference CDS (Biopython, match +2, mismatch −1, gap-open −2, gap-extend −0.5)^85^. Nucleotide substitutions and indels were called in reference coordinates. Amino-acid changes were called by grouping alignment columns into codons by reference position and translating each codon. Substitutions were partitioned into synonymous and nonsynonymous classes, and per-clone tables of NT and AA substitutions were compiled for downstream analysis. Supplementary Fig. 12C used Maple to generate de novo consensuses for comparison to clonal sequencing pre-refresh^5^. Lists of all nucleotide and amino acid substitutions presented in this work can be found in the Source Data file and raw sequencing files can be retrieved under the NCBI accession PRJNA1533496.

### Automated OrthoRep biosensor passaging

For the automated passaging campaign in Fig. 5, passages were initiated in two 24-well plates containing either loperamide (100 μM) or DMSO only. Automation-compatible plates were shaken at 250 rpm at 30 °C and OD600 was monitored continuously on a GrowthProfiler960 (Enzyscreen); when any well reached OD600 ≥ 3, all wells were passaged and normalized to a starting OD600 of 0.10 using the 96-channel variable-span head of a Dynamic Devices Lynx liquid handler. Plates were moved between the GrowthProfiler and the Lynx by a Thermo Spinnaker robotic arm, with the workflow orchestrated in Thermo Momentum. Culture density was tracked at each measurement by an XML “watchdog” script that parsed GrowthProfiler OD readings and triggered the passaging routine once the threshold was met. Manual interventions are noted in source data.

### High-throughput growth-based screening (HTS)

Growth-based selection screens were performed in 100 μL of biosensor media in 96-well plates, with overnight cultures back-diluted to an OD of ∼0.3 and agonist added at dilution. Plates were grown at 30 °C with 200 RPM shaking and growth (OD600) was recorded on a Tecan Sunrise plate reader. Agonist concentrations were chosen to impose selection near the wild-type response. For the Fig. 3D screen, cells were assayed at a limiting DAMGO concentration of 10 μM alongside a no-DAMGO control. For the OPRD1 screen in Fig. 5, cells were assayed at 50 μM (yielding approximately wild-type ON growth), 4 μM (limiting, to resolve variants outperforming wild type), and a no-agonist control.

### Drug-response curves

Agonist dilution series were prepared in DMSO as either log10, log2, or log3 dilutions depending on the range required (specified per figure legend), and an equal volume of DMSO was delivered to every concentration well and biological replicate so that solvent carryover was held constant across each series. Fluorescence-based response curves were acquired on a Cytek Aurora full spectrum flow cytometer. Samples were gated for debris and singlets, with a minimum of 2,000 events collected per sample after gating. Growth-based response curves were measured on a Tecan Infinite 200 pro as 48 h or 24 h endpoint readings. All curves were fit in python matplotlib using 4-parameter model to determine EC_50_ and the maximal response.

### Microscopy

For Supplementary Fig. 10, yeast cultured in synthetic media were mounted onto slides for brightfield imaging. Images were acquired on a DMi6000B microscope (Leica Microsystems) fitted with an HCX PL APO 63× objective, an Orca R2 CCD camera (Hamamatsu), and Volocity 5.5.1 software (PerkinElmer). Cell morphology was assessed qualitatively; no quantitative image analysis was performed. Images were assembled in Illustrator (Adobe).

### Statistics and reproducibility

No statistical method was used to predetermine sample size. Sample sizes were in line with generally accepted standards in the field. No data were excluded from the analyses. The experiments were not randomized, and investigators were not blinded to allocation during experiments or outcome assessment. Statistical analyses and curve fitting were performed in GraphPad Prism 8, Microsoft Excel 16, or using custom Python 3.14 scripts; details of the tests applied are given in the relevant figure legends.

