## Supplementary Figures for "Heterologous engineering of receptors using OrthoRep (HERO) for directed evolution of GPCRs in yeast"

### Dykstra *et al.* Supplementary figures

#### Contents

1. Development of the OrthoRep biosensor strain
2. Protoplast fusion and ploidy validation of OrthoRep biosensor strains
3. HIS3 reporter modulation with 3-AT
4. High-stringency passaging with 3-AT
5. FACS OPRM1 library sorting plots
6. Early DAMGO-selected mutation profile of OPRM1 from 611-DNAP1 biosensor
7. Endpoint DAMGO- and loperamide-selected mutation profile of OPRM1 from 611-DNAP1 biosensor
8. Unsorted OPRM1 biosensor after 14 DAMGO-selected passages from 611-DNAP1 biosensor
9. Ultra-low stringency passaging (selection escape)
10. Escaper phenotype
11. Duplicated GPA1-GNAI3 copy ablates signalling in the growth-based biosensor
12. Individual amino acid substitution overlap pre- and post- background refresh
13. High-stringency passaging with 3-AT after background refresh
14. Mutation frequency of OPRM1 after 14 + 6 passages
15. Post refresh library growth under ligand-dependent growth and non-selective conditions
16. Histograms of ENVY induction from plasmid-based expression of WT and Mutant OPRM1 with ergosterol/cholesterol
17. Additional serial-passaging growth phenotypes of OPRM1 in the HERO strain
18. Mutational frequency across the BadBoy3 evolution campaign
19. Continuous directed mutagenesis of human OPRD1 receptor against SNC-80
20. Mutational frequency of OPRD1 increase from passage 5 to passage 10
21. Well-level resolution of mutational frequency of OPRD1 during automated evolution in the presence of loperamide
22. Single-copy OPRD1 responds to loperamide and SNC-80
23. Additional isolated mutants from OPRD1 high throughput evolution

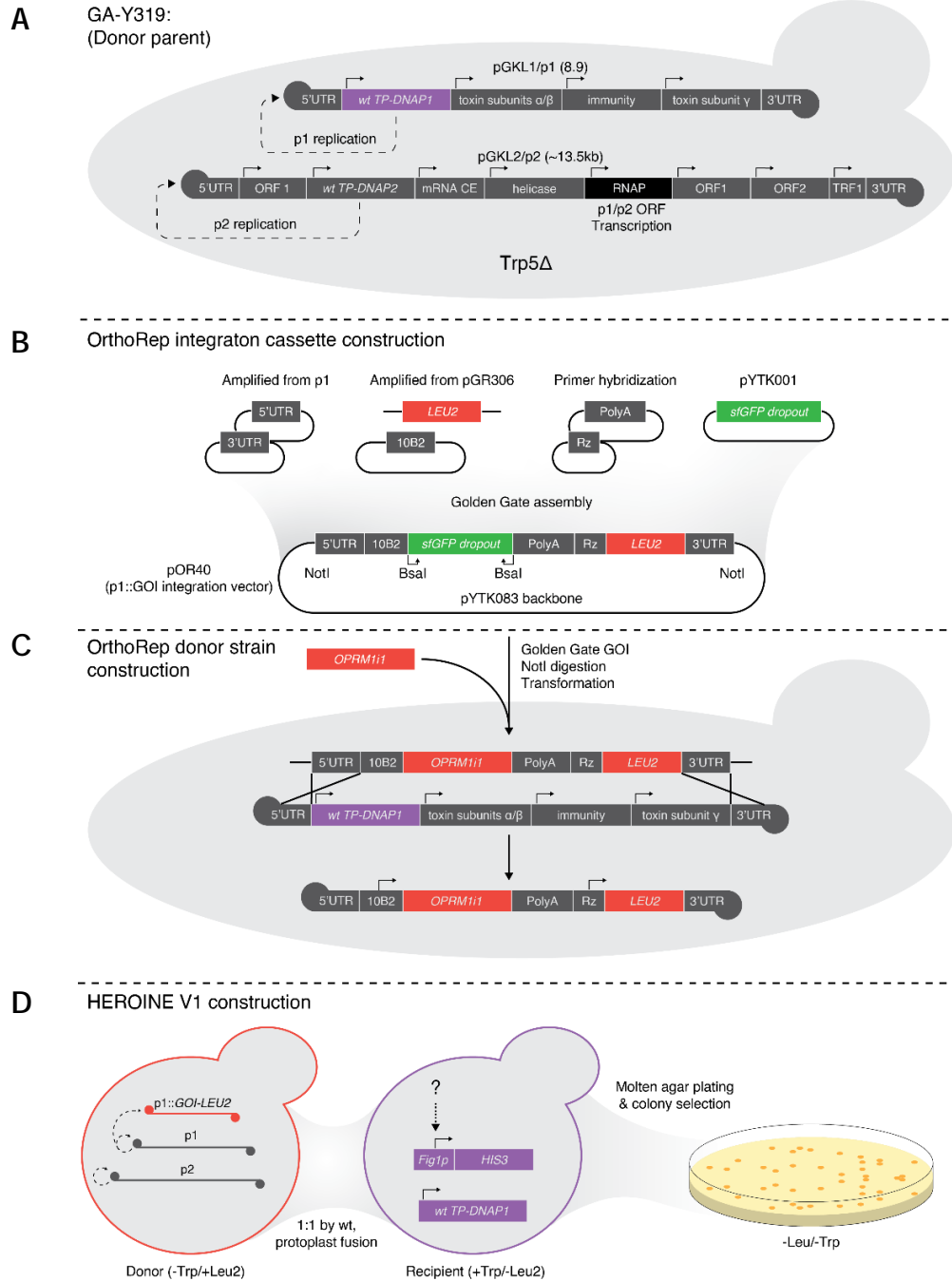

**Supplementary Fig. 1. Development of the OrthoRep Biosensor strain.** **A** GA-Y319 donor parental strain and wild-type linear, cytosolic p1/p2.<sup>1</sup> **B** p1 integration cassette Golden Gate–assembled from level-0 part plasmids or amplicons<sup>2</sup> **C** Golden Gate insertion of OPRM1 (GOI) and linear-donor transformation into GA-Y319; homology-directed repair inserts the cassette into p1, and wild-type p1/p2 maintain p1::GOI-LEU2 copy number, which in turn maintains selection in –Leu. **D** Protoplast fusions of the resulting donor and a recipient bearing TP-DNAP1 *in trans*. Recipient colonies (Trp1) that are Leu2 received p1::GOI-LEU2 and the cytosolic components necessary for replication and transcription.

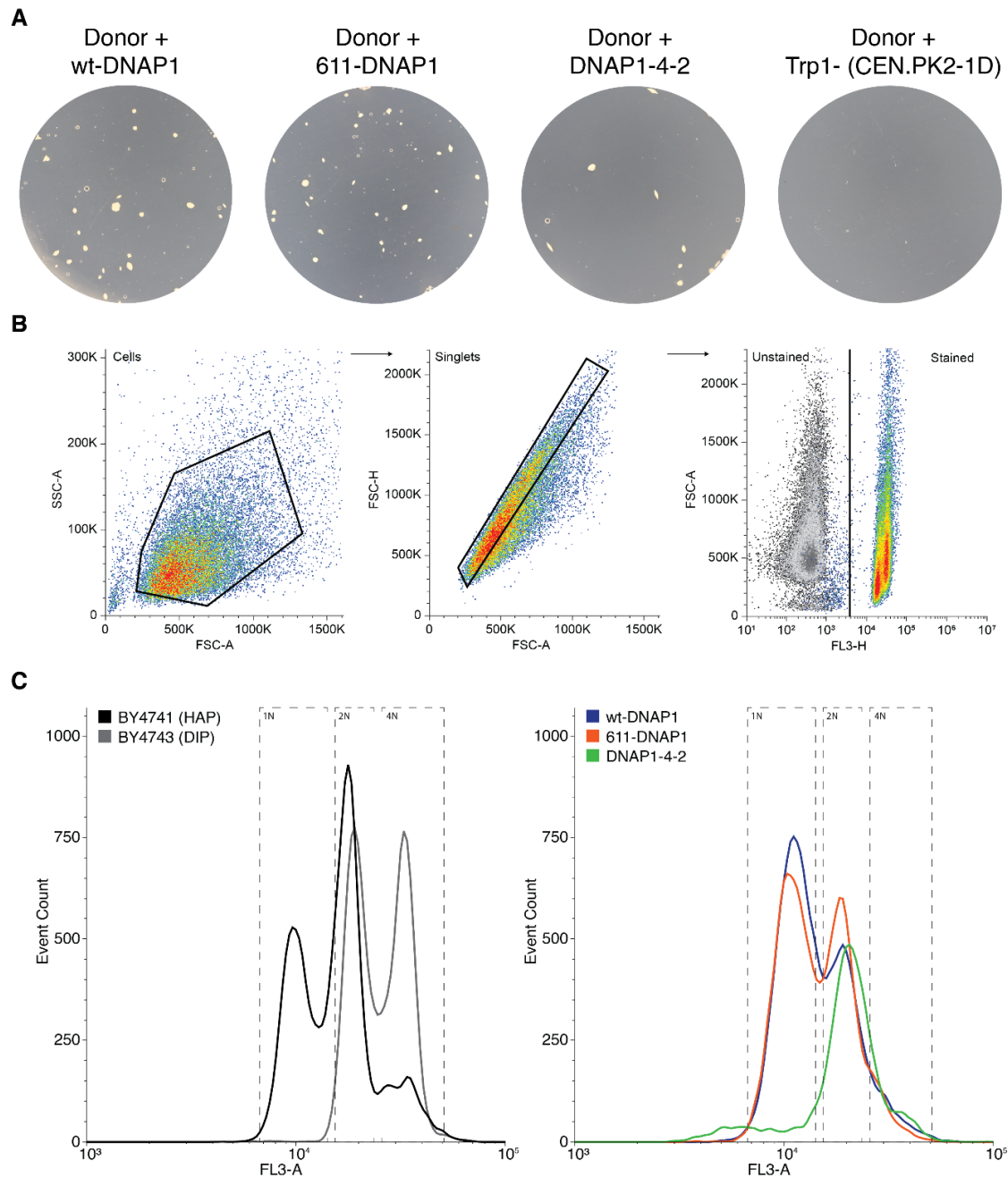

**Supplementary Fig. 2. Protoplast fusion and ploidy validation of OrthoRep biosensor strains. A** Agar-embedded protoplast fusants generated by fusing a donor harboring $p1::OPRM1-LEU2$ with recipients carrying wt-DNAP1 (non-mutagenic), 611-DNAP1 (mid-mutagenic), or DNAP1-4-2 (highly mutagenic), alongside a $trp^-$ recipient control. **B** Representative gating strategy used for propidium iodide staining and subsequent flow cytometry experiments; samples were gated to exclude debris and doublets, and fluorescent populations were identified by gating against unstained controls. **C** Propidium iodide histograms of representative fusants from (A). Gates for 1N, 2N, and 4N peaks were defined using haploid (BY4741) and diploid (BY4743) controls and applied to fusants.

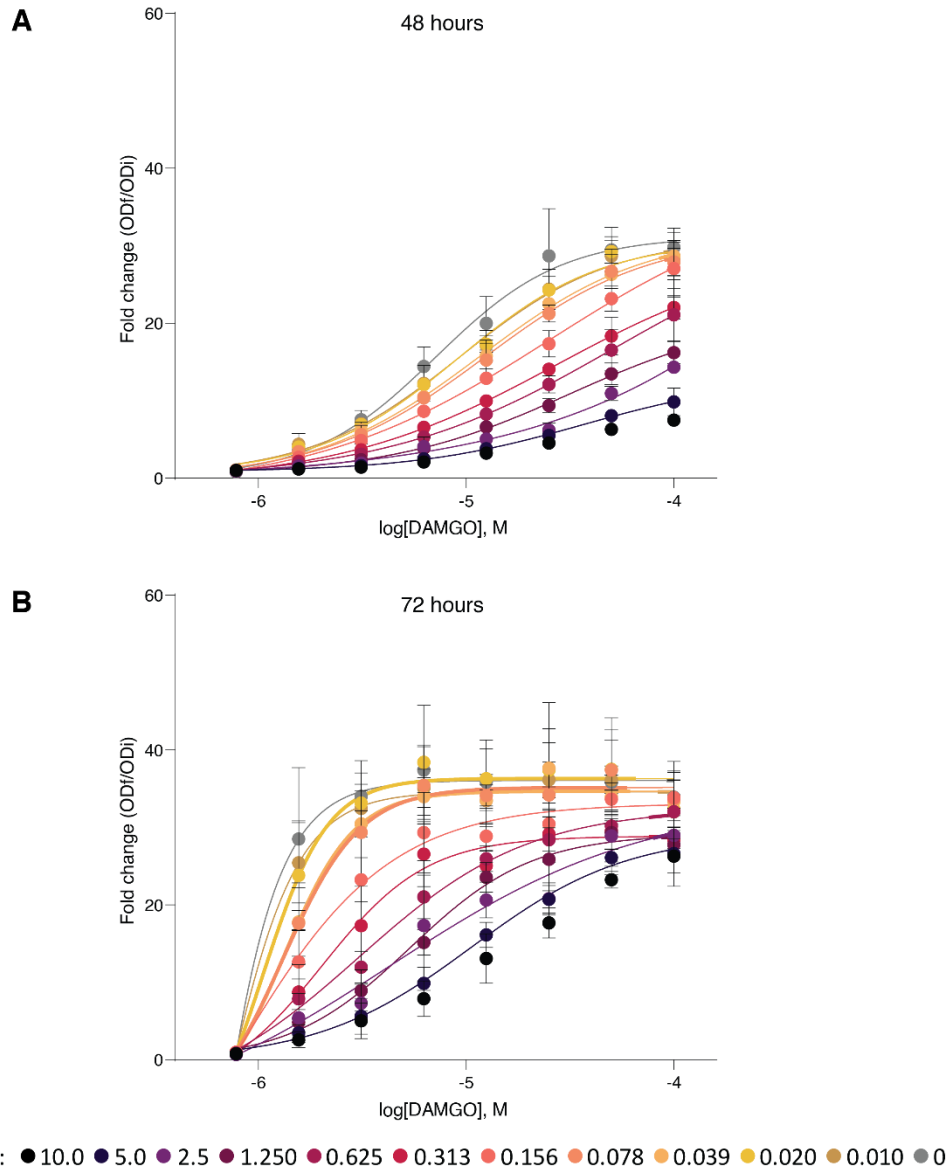

**Supplementary Fig. 3. HIS3 reporter modulation with 3-AT.** Dose–response of the wt-DNAP1 biosensor strain ($p1::OPRM1$-$LEU2$, $wt$-$DNAP1$ in trans) to DAMGO across a two-fold dilution series of 3-AT (0–10 mM), measured as fold change in optical density (ODf/ODi) after **A** 48 h and **B** 72 h of growth. Increasing 3-AT progressively suppressed basal HIS3 read-through and shifted the apparent dynamic range of the DAMGO response, with the effect most pronounced at 72 h, where lower 3-AT concentrations approached saturation. At 3-AT concentrations $\geq 2.5$ mM the DAMGO fold-change did not saturate, leading us to adopt 2.5 mM for subsequent high-stringency evolution campaigns. Data are mean ± s.d. of three biological replicates.

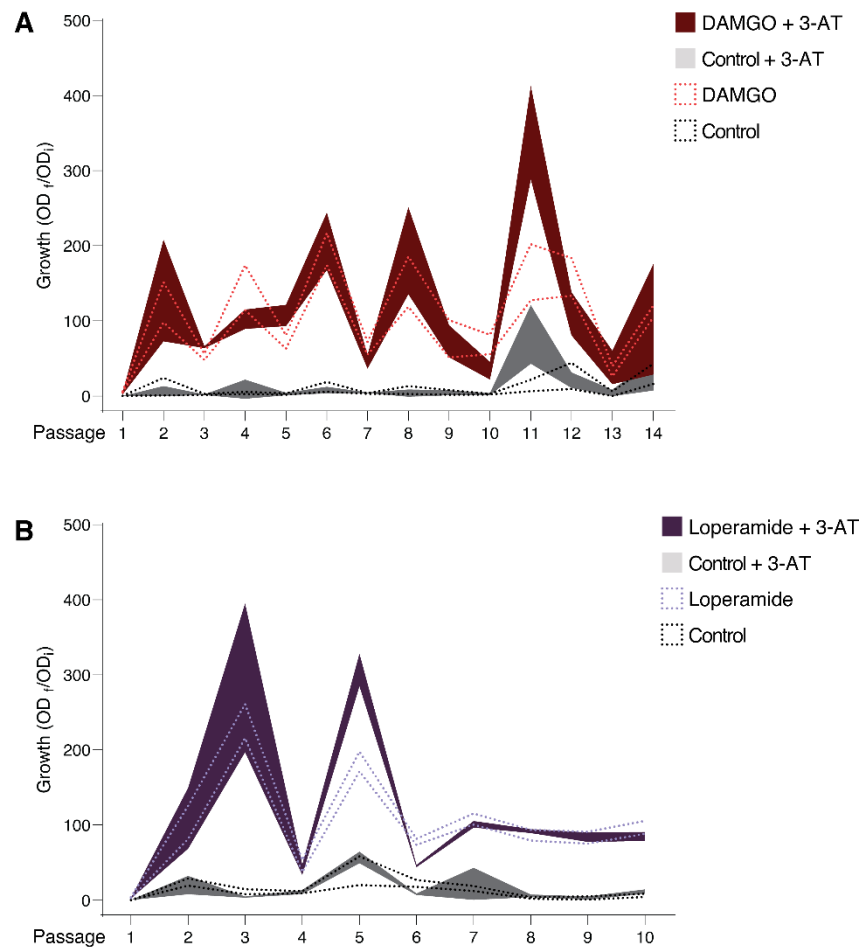

**Supplementary Fig. 4. High-stringency passaging with 3-AT.** OrthoRep biosensor with p1::*OPRM1-LEU2* strain growth with 3-AT supplemented with **A** DAMGO or **B** loperamide (50 μM each), shown as solid fill. Dotted curves are low-stringency passages from Fig. 2A,B shown for comparison. Curve area represents mean ± s.d. growth fold change.

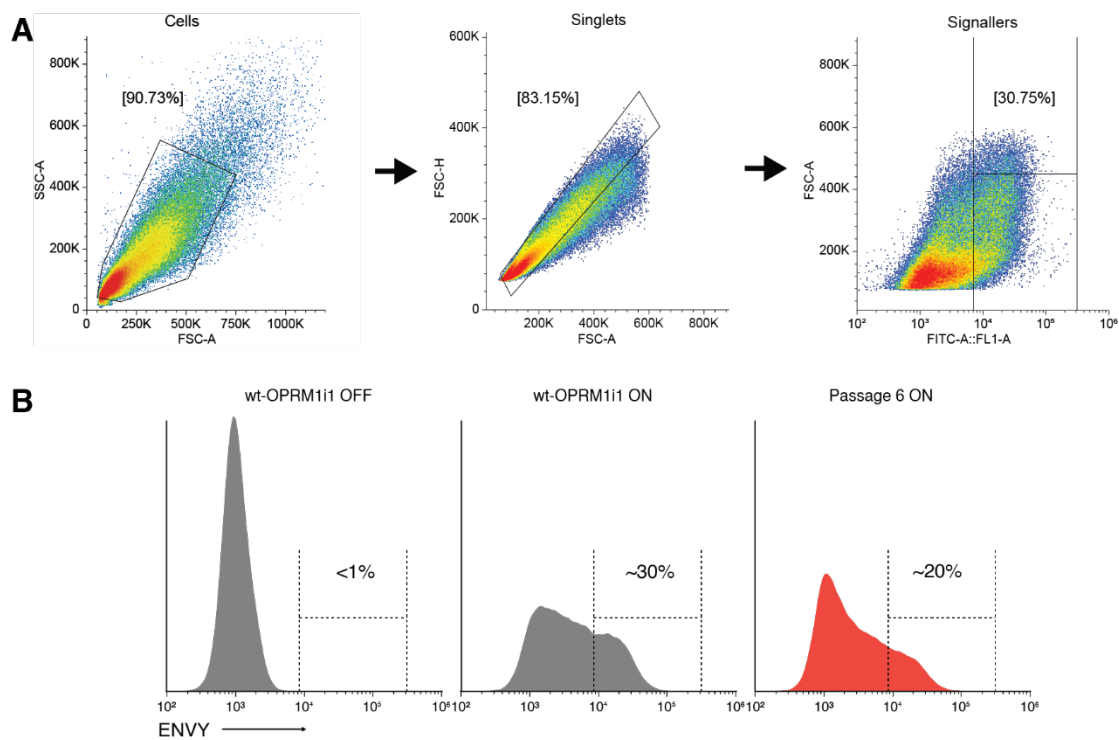

**Supplementary Fig. 5. FACS OPRM1 library sorting plots.** **A** FACS gating strategy of *wt-OPRM1* control transformed into a fluorescent biosensor. Cells are gated to remove debris and doublets. Each experiment sorted using a gate representing the top 30% of *wt-OPRM1* signallers as a function of ENVY fluorescence. **B** Histograms of ENVY expression of the *wt-OPRM1* without DAMGO, *wt-OPRM1* with 50 μM DAMGO, and the DAMGO-evolved (passage 6) library with 50 μM DAMGO.

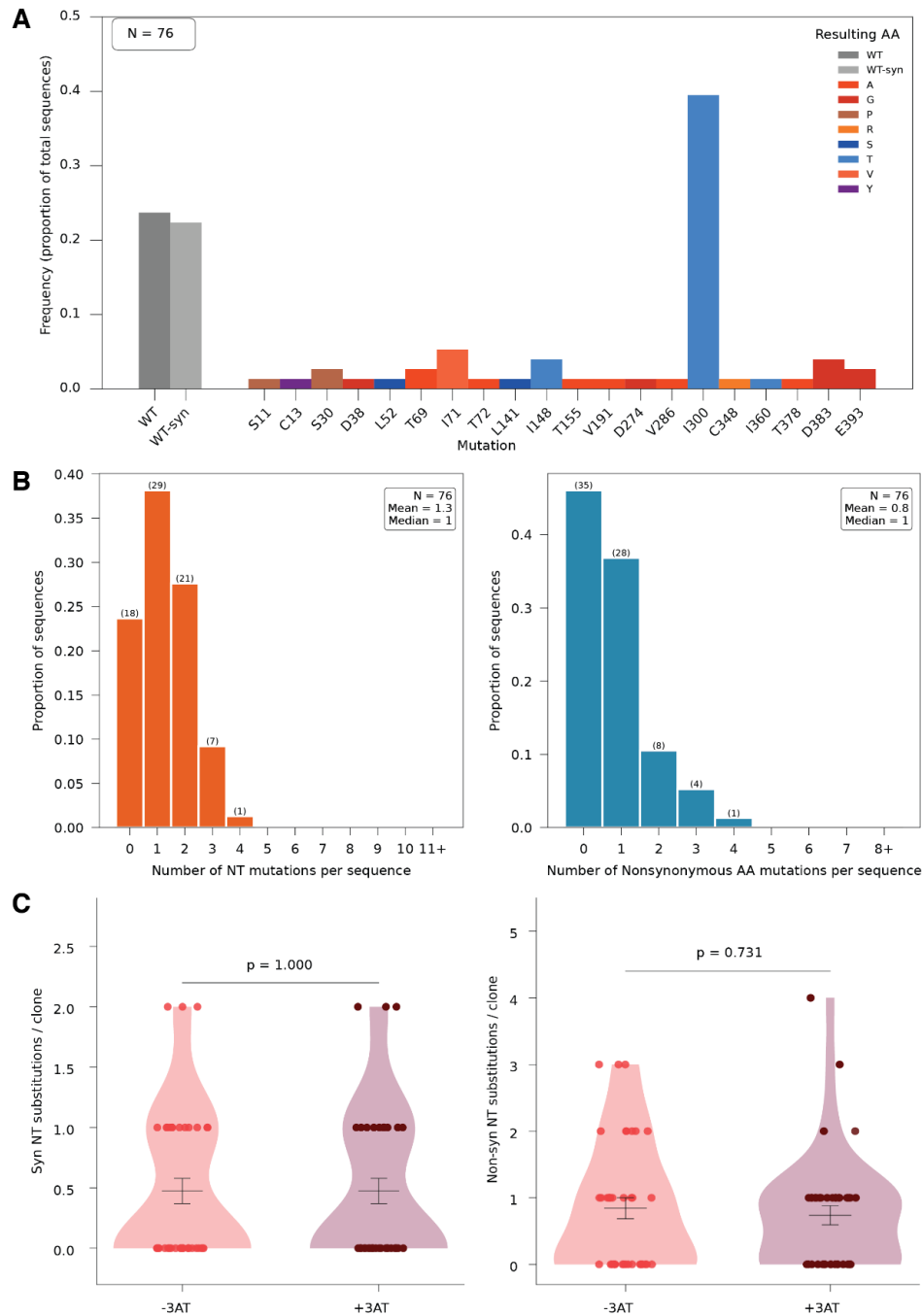

**Supplementary Fig. 6. Early DAMGO-selected mutation profile of OPRM1 from 611-DNAP1 biosensor.** Populations were passaged six rounds in 50 μM DAMGO with or without 3-AT (2.5 mM), then sequenced from genomically integrated fluorescent biosensor clones. **A** Per-position amino-acid mutation frequencies across OPRM1. **B** Per-clone mutation frequency for nucleotide (orange) and non-synonymous amino-acid (blue) substitutions (aggregate of +3-AT and -3-AT groups; mean = 1.3 NT substitutions/clone). **C** Synonymous and non-synonymous nucleotide substitutions per clone, split by ±3-AT condition (two-sided Mann–Whitney U). Data in **A** and **B** aggregate the +3-AT and -3-AT groups.

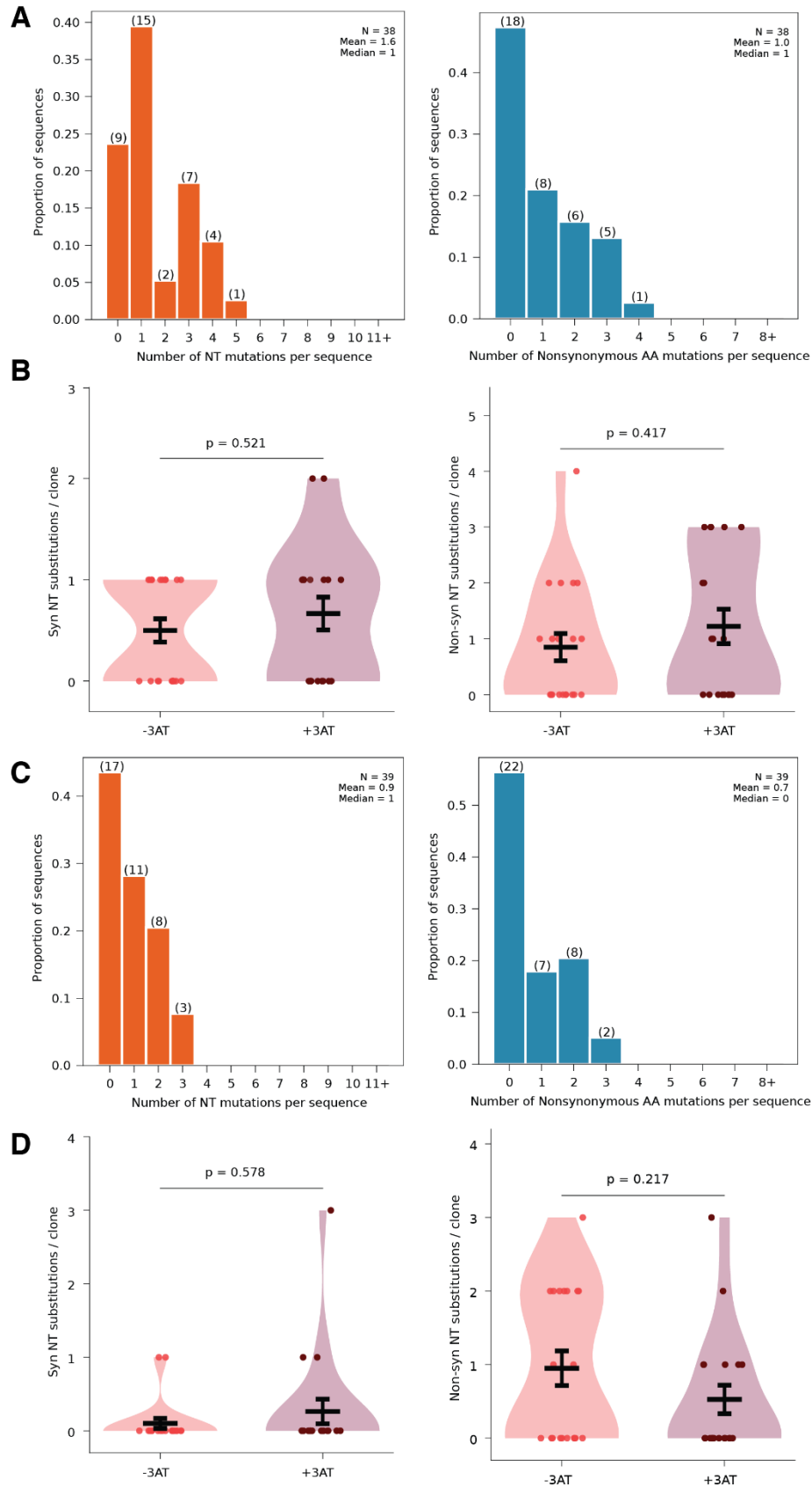

**Supplementary Fig. 7. Endpoint DAMGO- and loperamide-selected mutation profile of OPRM1 from 611-DNAP1 biosensor.** Populations were passaged 14 rounds in 50 μM DAMGO or 10 rounds in 50 μM of loperamide with or without 3-AT (2.5 mM), then sequenced from FACS-sorted, genomically integrated fluorescent biosensor clones. **A** DAMGO Per-clone mutation frequency for nucleotide (orange) and non-synonymous amino-acid (blue) substitutions (aggregate of +3-AT and -3-AT groups; mean = 1.6 NT substitutions/clone). **B** DAMGO Synonymous and non-synonymous nucleotide substitutions per clone, split by ±3-AT condition (two-sided Mann–Whitney U). **C** Loperamide per-clone mutation frequency for nucleotide (orange) and non-synonymous amino-acid (blue) substitutions (aggregate of +3-AT and -3-AT groups; mean = 0.9 NT substitutions/clone). **D** Loperamide Synonymous and non-synonymous nucleotide substitutions per clone, split by ±3-AT condition (two-sided Mann–Whitney U). Data in A and C aggregate the +3-AT and -3-AT groups.

**A**

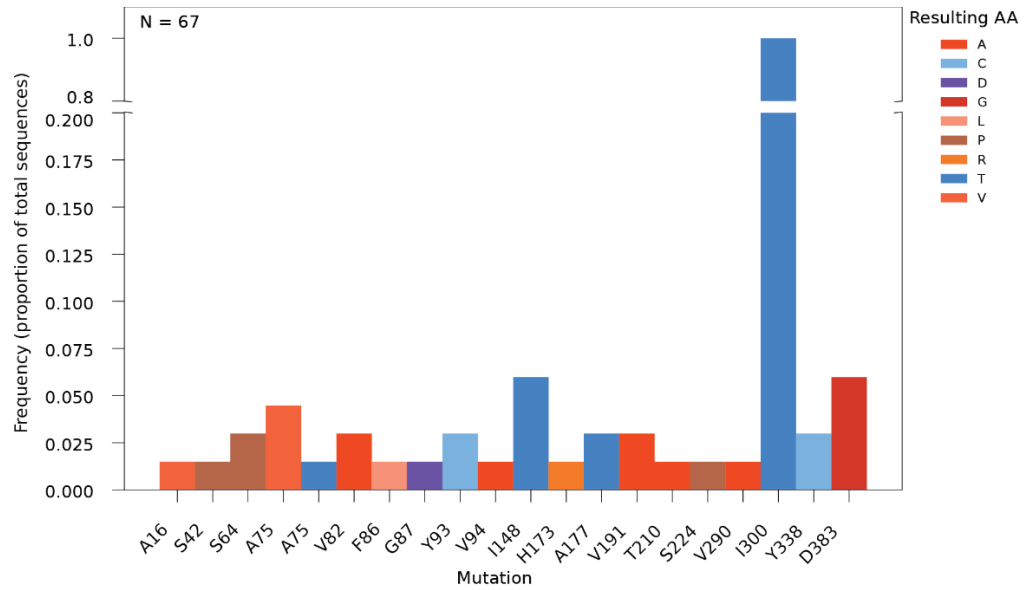

**B**

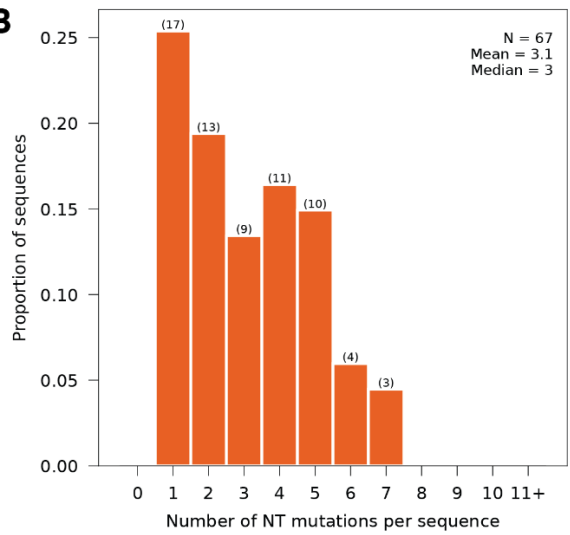

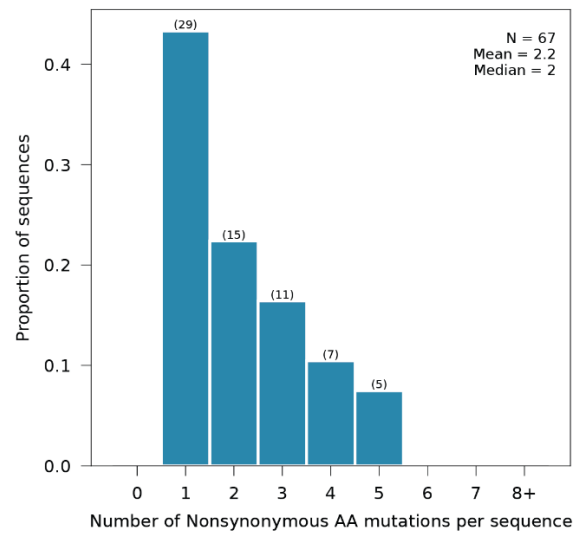

**C**

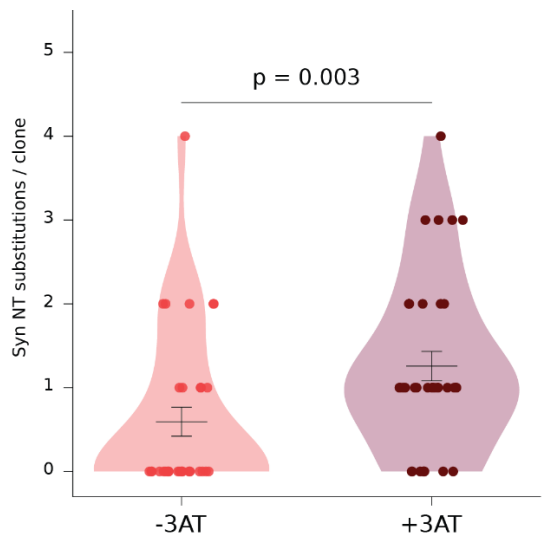

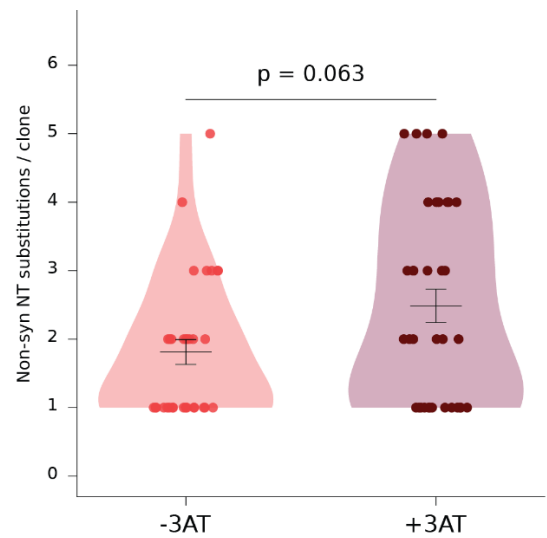

**Supplementary Fig. 8. Unsorted OPRM1 biosensor after 14 DAMGO-selected passages from 611-DNAP1 biosensor.** **A** Per-position amino-acid mutation frequencies across *OPRM1* after 14 rounds of passages and randomly picked without sorting. **B** NT (orange) and AA (blue) substitutions after passaging 611-DNAP1 biosensor bearing p1::*OPRM1-LEU2* for 14 rounds in 50 μM DAMGO. **C** Synonymous and non-synonymous nucleotide substitutions per clone, split by ±3-AT condition (two-sided Mann–Whitney U). Data in **A** and **B** aggregate the ±3-AT conditions.

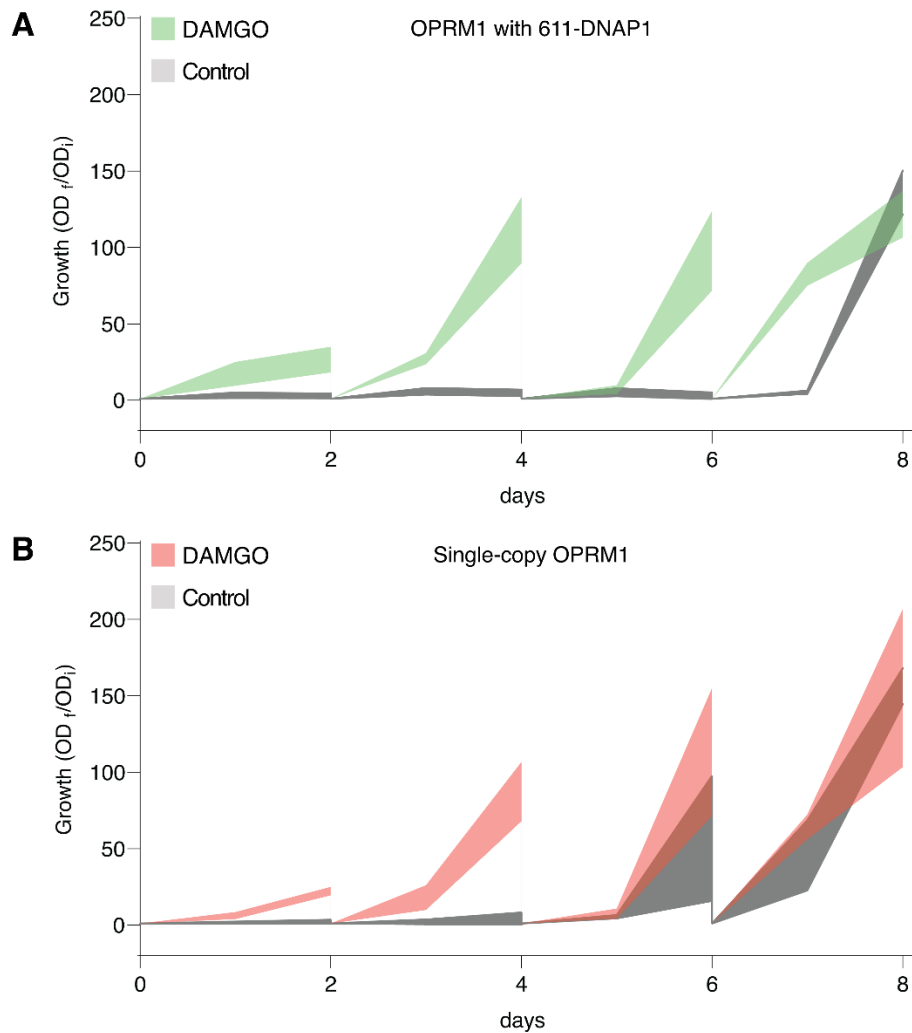

**Supplementary Fig. 9. Ultra-low stringency passaging (selection escape).** **A** 611-DNAP1 strain bearing $p1::OPRM1-LEU2$ and **B** single-copy, integrated OPRM1 biosensor grown in 2 mL SC-His supplemented with 50 μM DAMGO or control (DMSO) and back-diluted 1:100 every 48 h ($n = 3$ biological replicates). At each passage, only DAMGO-treated replicates were carried forward into fresh drug or control medium; controls report the rate of selection escape. Shaded bands span mean ± s.d. of the replicate fold changes. In this campaign, where $p1::OPRM1-LEU2$ was maintained by downstream reporter activation alone ($pFIG1-HIS3$), selection escape occurred after four passages ($\sim 25$ generations) in both strains.

**A No escape**

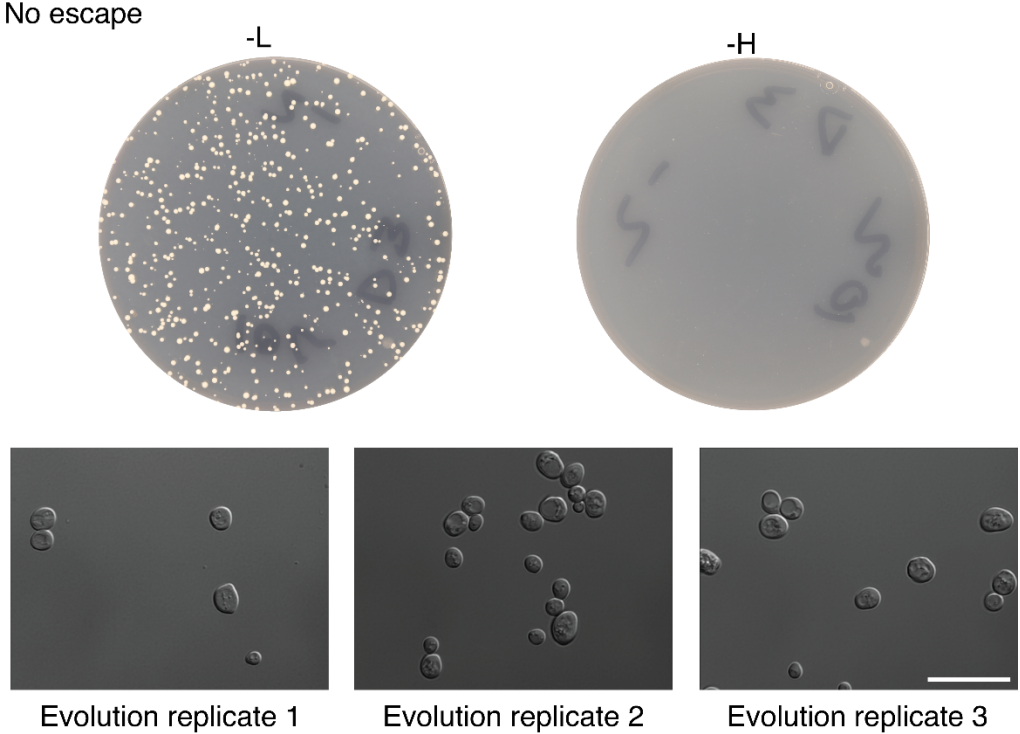

**B Escape**

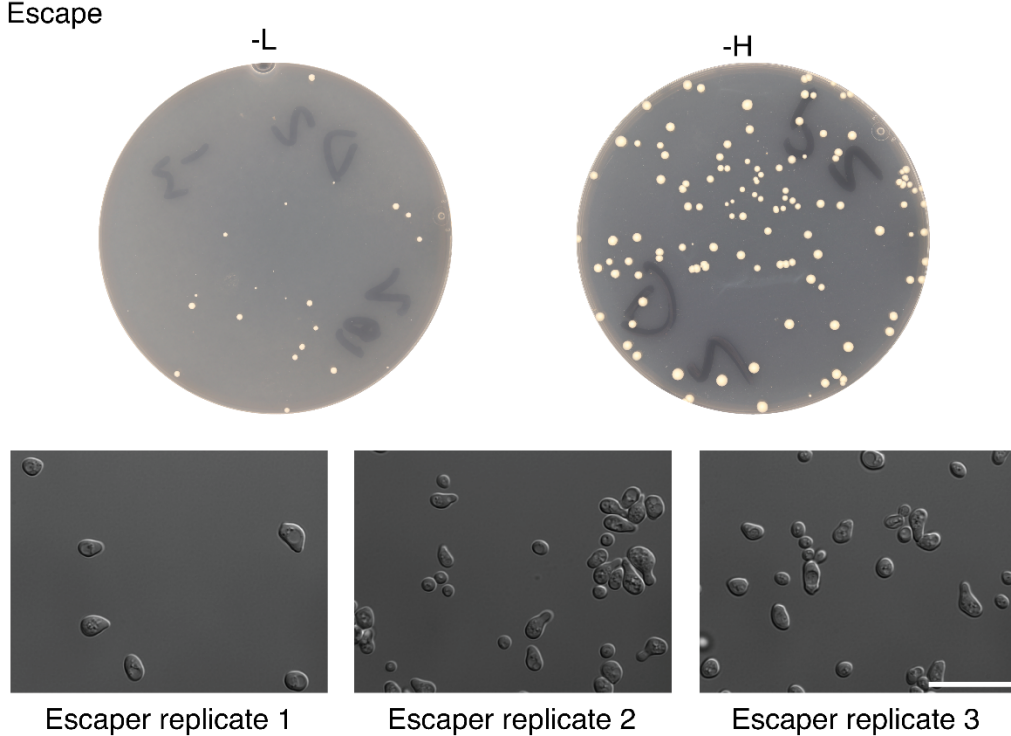

**Supplementary Fig. 10. Escaper phenotype.** Representative colony growth on SC-Leu (p1::*OPRM1-LEU2*) and SC-His media, with yeast micrographs, for non-escape **A** and escaped **B** populations across three evolution replicates. Scale bar represents 10 μm. Sequencing detected mutations of *GPA1-GNAI3* in 2/3 clones from B (Source Data), indicating that the escape phenotype arises from host-encoded *GPA1* loss-of-function rather than p1-mediated mutagenesis of the receptor cassette.

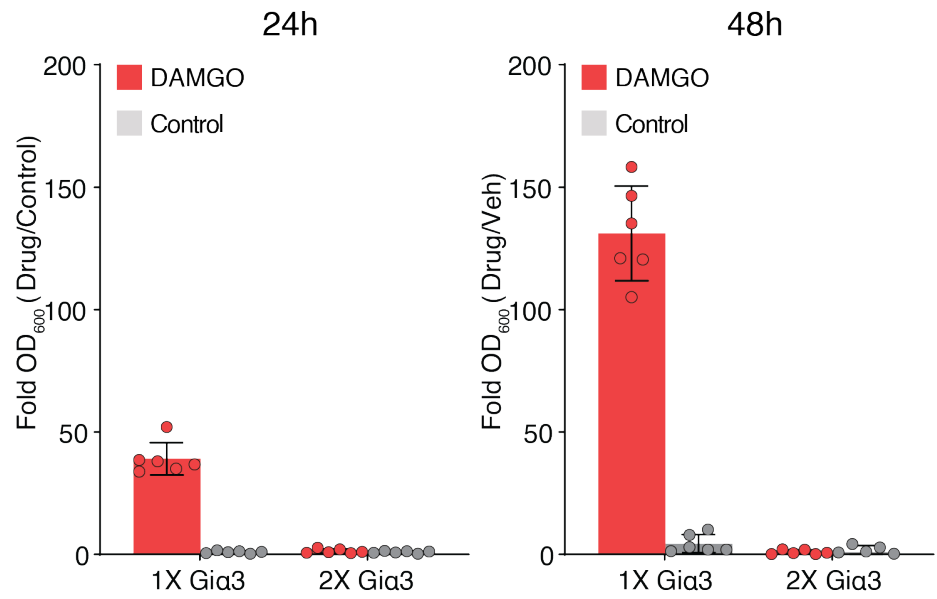

**Supplementary Fig. 11. Duplicated *GPA1-GNAI3* copy ablates signalling in the growth-based biosensor.** Integrating a second copy of the GPA1-GNAI3 (Gαi3) chimera at a safe-harbor locus (FF20) ablated ligand-dependent signalling. Bars show growth fold change relative to the mean OD<sub>600</sub> of each group's control replicates at 24 and 48 h, plotted as mean ± s.d. (n = 6).

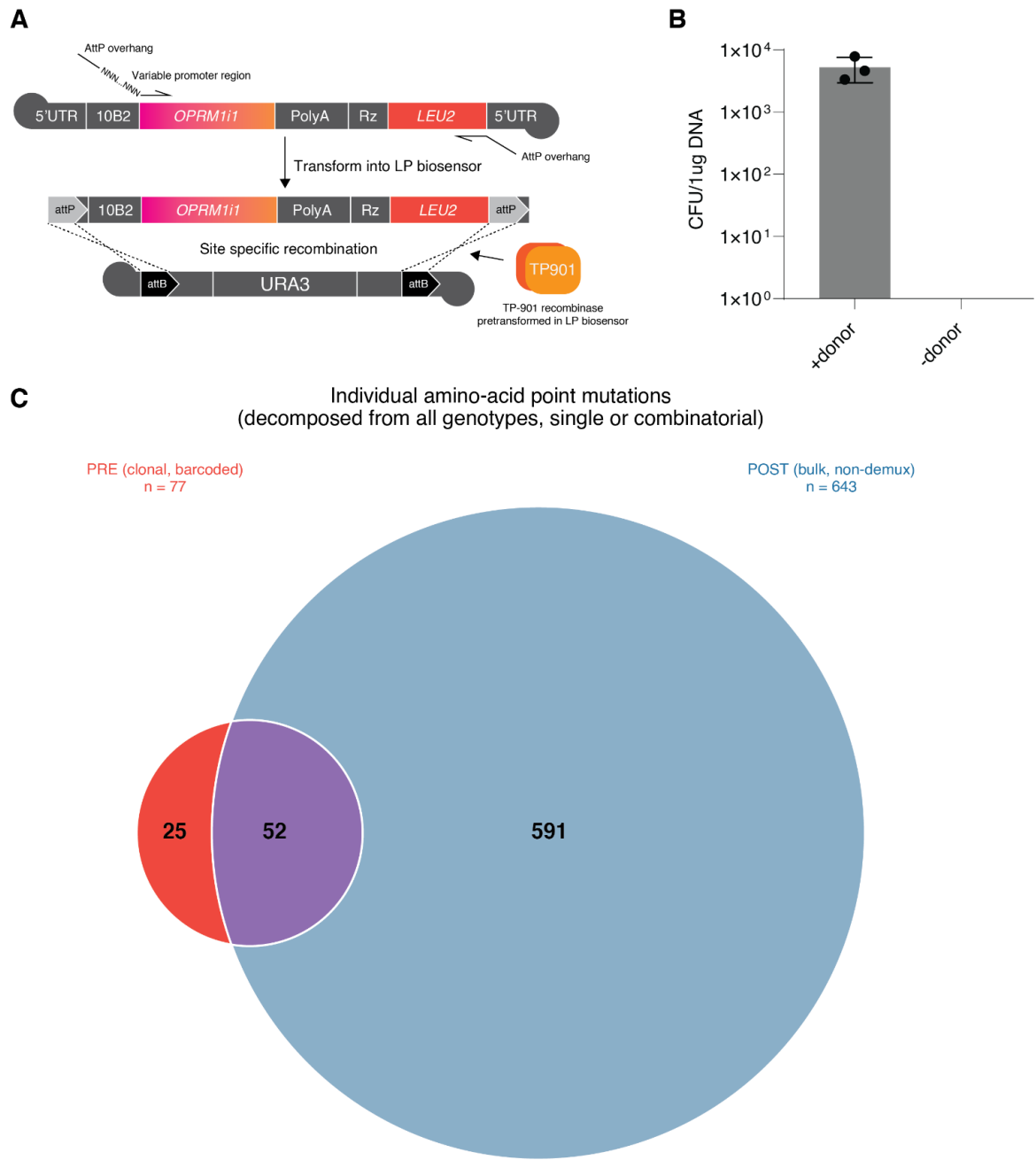

**Supplementary Fig. 12. Individual amino acid substitution overlap pre- and post- background refresh.** **A** TP901 integrase-mediated library refreshing onto the p1 landing pad. **B** Colony-forming units per $\mu\text{g}$ of donor DNA from passage 14 DAMGO-selected variants transformed into the 611-DNAP1 landing strain; a quarter of the reaction was plated on SC-Leu (recombinant p1 marker) and normalized to the amount of DNA provided ($n = 3$). **C** Amplicons from boiled cultures of DAMGO and DAMGO +3-AT, passage 14 were pooled and transformed directly without purification. After refreshing the background, recovering in non-biosensor-selective medium, and passaging 48 h in 50 $\mu\text{M}$ DAMGO (passage 14 + 1) cultures were pooled and bulk sequenced. Sequences were assembled with minimap2/Medaka to generate unique genotypes, then compared to concatenated amino-acid substitutions from the DAMGO 50 $\mu\text{M}$ passage-14 analysis.

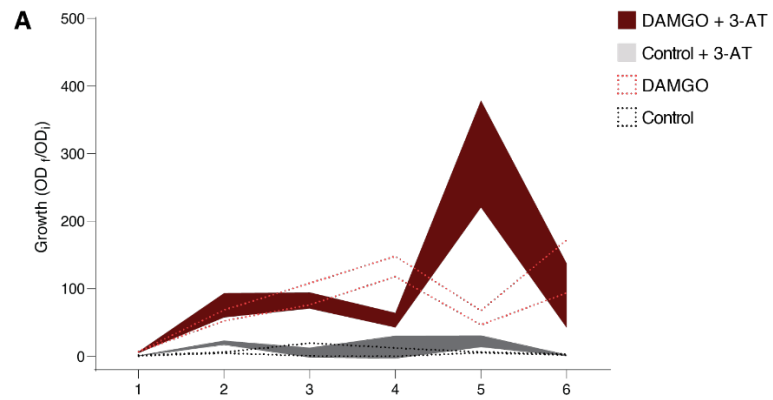

**Supplementary Fig. 13. High-stringency passaging with 3-AT after background refresh.** OrthoRep landing pad p1 strain with pre-evolved OPRM1-LEU2 growth with 3-AT supplemented with DAMGO 50 μM shown as solid fill. Dotted curves are low-stringency passages without 3-AT from Fig. 3B shown for comparison. Curve area represents mean ± s.d. growth fold change.

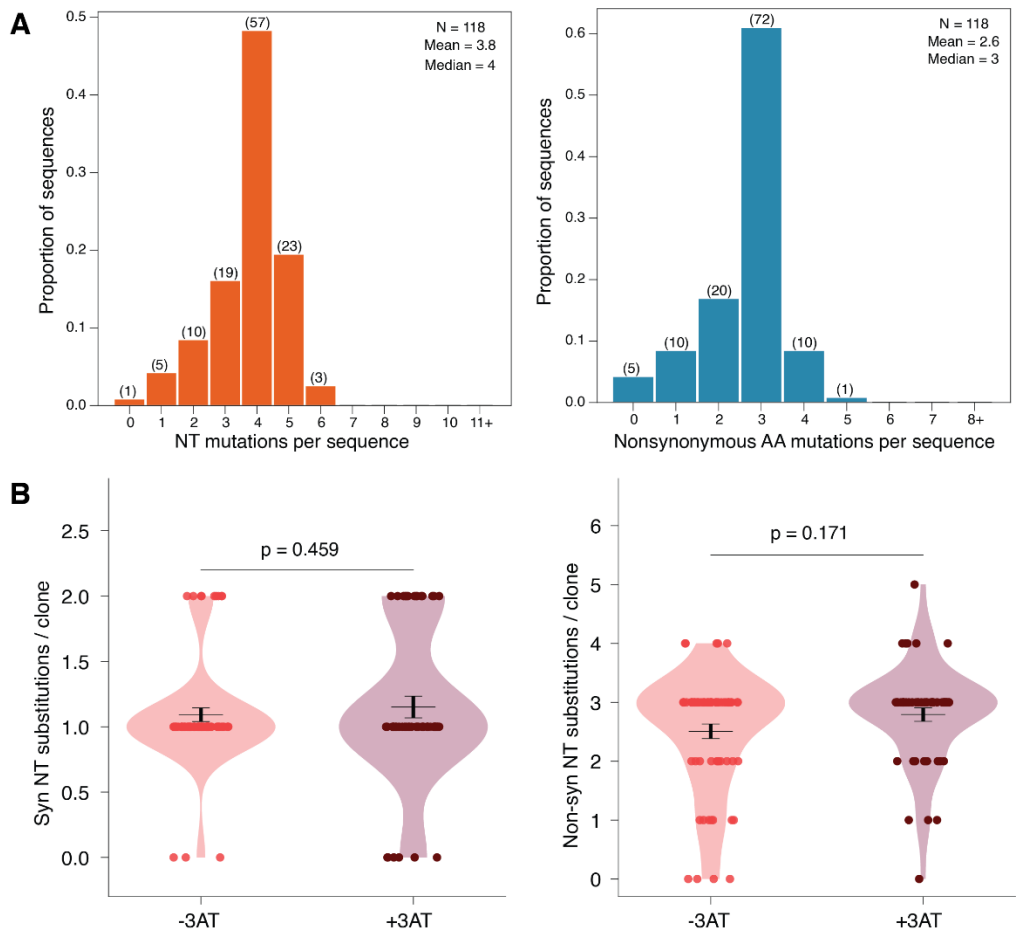

**Supplementary Fig. 14. Mutation frequency of OPRM1 after 14 + 6 passages.** Human μ-opioid receptor variant OPRM1 evolved with DAMGO (50 μM), passaged 14 times, refreshed into a fresh host background, and passaged a further 6 times before individual colonies were picked at random (n = 118). **A** Aggregate mutation frequency across the selected pool: left, nucleotide (NT) substitutions per sequence (mean = 3.8, median = 4); right, non-synonymous amino-acid substitutions per sequence (mean = 2.6, median = 3). Bars show the proportion of sequences in each bin; parenthetical values give the underlying sequence counts. **B** Per-clone substitution frequency partitioned by whether the final 6 passages were under *HIS3* selection (+3-AT) or without (-3-AT): left, synonymous NT substitutions per clone; right, non-synonymous NT substitutions per clone (-3-AT, n = 65; +3-AT, n = 53). Violin plots show the full distribution with overlaid individual clones; horizontal bars and whiskers denote mean ± s.d. Neither the synonymous (p = 0.459) nor the non-synonymous (p = 0.171) frequency differed significantly between conditions (two-sided Mann-Whitney U), indicating that the added 3-AT selection did not detectably alter overall mutational frequency in this campaign.

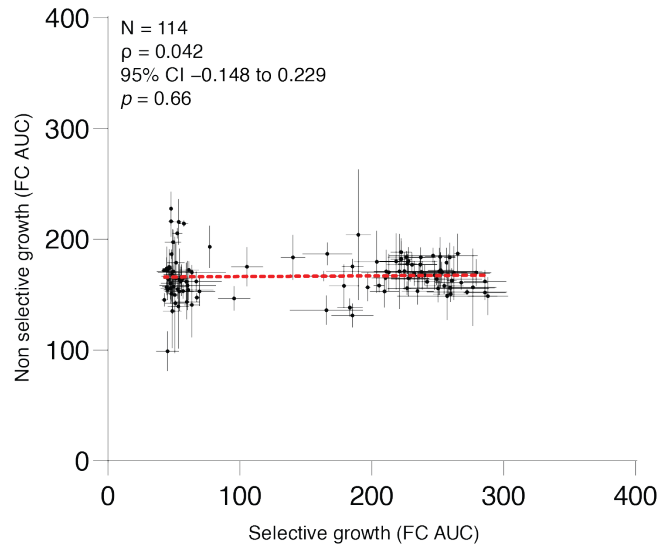

**Supplementary Fig. 15. Post refresh library growth under ligand-dependent growth and non-selective conditions.** Selective (ligand-dependent) versus non-selective growth for individual mutants from the refreshed library in the single-copy sensor background ($n = 114$). Selective growth was measured as fold-change AUC under 10 $\mu\text{M}$ DAMGO selection ($-\text{His}$), and non-selective (SC complete) growth as fold-change AUC in synthetic complete media. Each point is a single mutant clone ($n = 3$); error bars denote s.d. on each axis. The red dashed line is a linear fit shown as a guide to the eye. Signalling-dependent and non-selective growth are uncorrelated (Spearman $\rho = 0.042$, $p = 0.66$, 95% CI $-0.148$ to $0.229$).

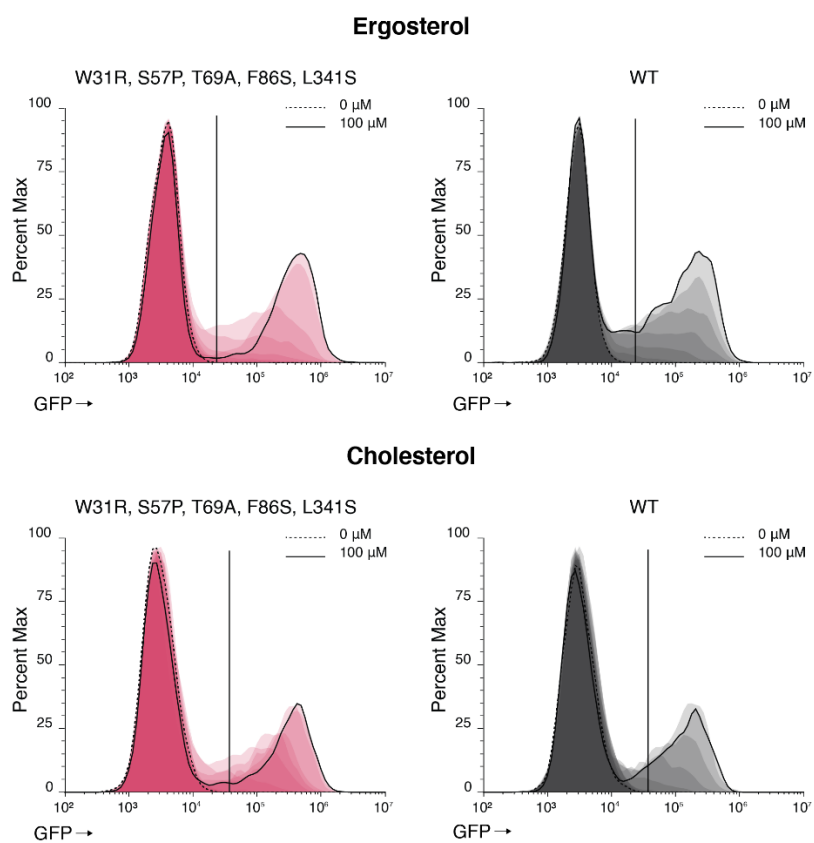

**Supplementary Fig. 16. Histograms of ENVY induction from plasmid-based expression of WT and Mutant OPRM1 with ergosterol/cholesterol.** Representative flow-cytometry histograms of ENVY fluorescence (biosensor output) for the evolved OPRM1 variant (W31R, S57P, T69A, F86S, L341S; pink) and wild-type OPRM1 (grey), expressed from a plasmid in yeast engineered to produce either ergosterol (top) or cholesterol (bottom) as the dominant membrane sterol (cf. Fig. 3E,F). Cells were treated with a 12-point three-fold DAMGO dilution series from control (0 μM; dashed) to 100 μM (solid); the vertical line marks the ENVY+ gating threshold.

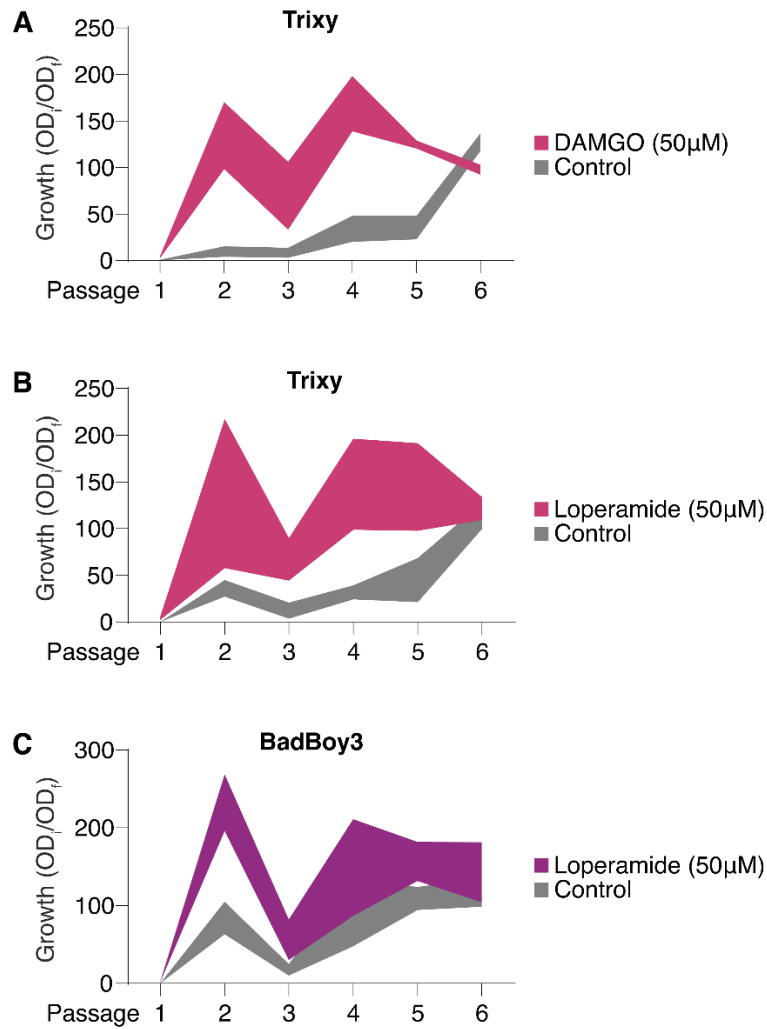

**Supplementary Fig. 17. Additional serial-passaging growth phenotypes of OPRM1 in the HERO strain.** Complementing the sequenced BadBoy3/DAMGO series in Fig. 4G, OPRM1 was introduced onto p1 of the Trixy and BadBoy3 HERO strains and serially passaged for six rounds in medium containing either an agonist or control; growth at each passage is fold change in culture density ($OD_i/OD_1$). The gap between agonist and control reflects ligand-dependent growth. Across all conditions, cultures remained viable and expandable over six passages, showing that the strain tolerates repeated propagation. **A** Trixy with DAMGO (50 µM) versus control. **B** Trixy with loperamide (50 µM) versus control. **C** BadBoy3 with loperamide (50 µM) versus control. Isolates from these supplementary series were not sequenced.

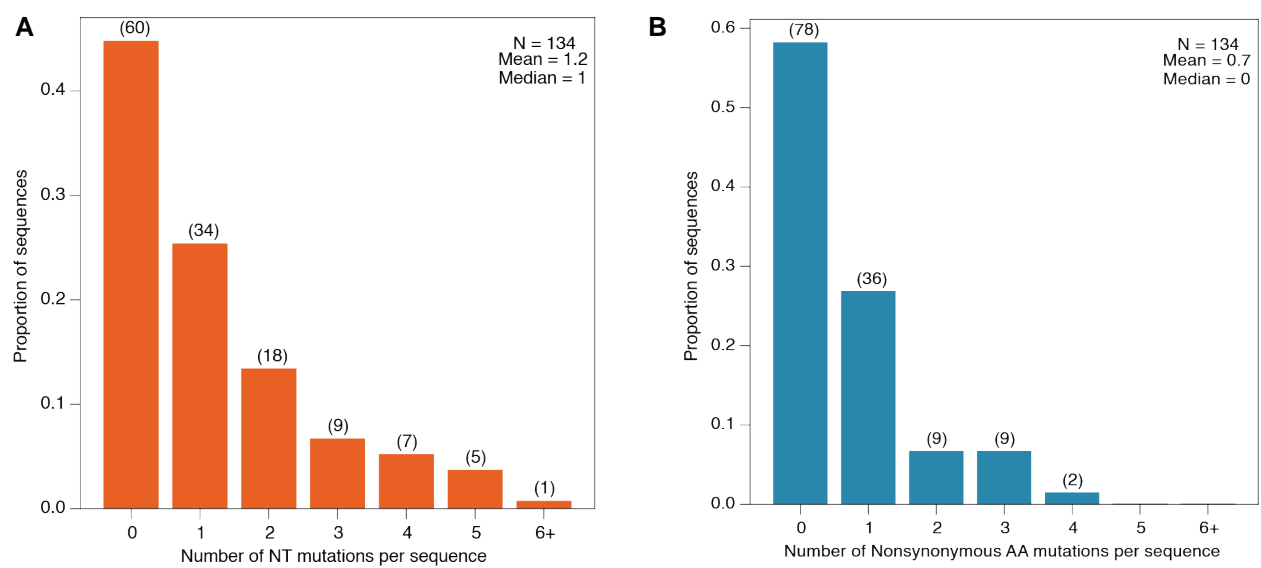

**Supplementary Fig. 18. Mutational frequency across the BadBoy3 evolution campaign.** Distribution of per-sequence mutation counts for clones recovered from the BadBoy3 OPRM1 evolution series (n = 134 sequences). Bars show the proportion of sequences carrying a given number of mutations; the raw sequence count in each bin is given in parentheses, with summary statistics inset. **A** Nucleotide (NT) mutations per sequence relative to the reference (mean 1.2, median 1). **B** Non-synonymous amino-acid (AA) mutations per sequence (mean 0.7, median 0). Both distributions are dominated by low-frequency sequences, the modal class is zero in each case, with a long tail of more heavily mutated clones, consistent with the low, distributed mutagenesis expected from the OrthoRep-driven campaign.

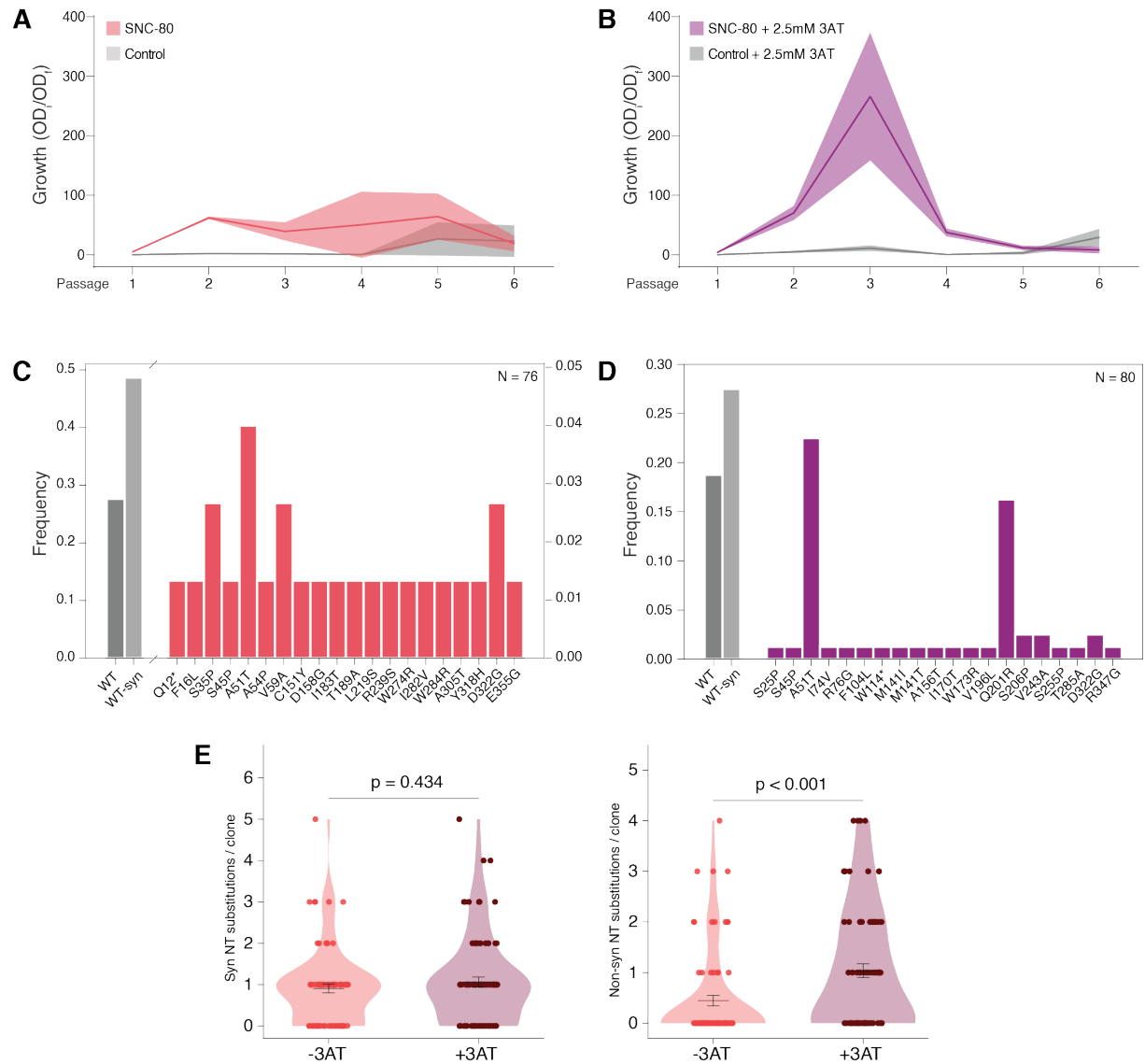

**Supplementary Fig. 19. Continuous directed mutagenesis of OPRD1 against SNC-80.** **A, B** Enrichment of SNC-80-responsive human δ-OR (OPRD1) variants over six serial passages of continuous evolution, measured as growth (OD<sub>i</sub>/OD<sub>i</sub>) at each passage. Cultures were selected with the δ-opioid-selective agonist SNC-80, 5 μM, or control, either without added 3-AT (3-amino-1,2,4-triazole) (A) or with 2.5 mM 3-AT to raise *HIS3* selection stringency (B). Solid lines indicate the mean and shaded regions ± s.d. across four replicate lineages. **C, D** Frequency of amino-acid substitutions recovered by transformation into CDY104 and sequencing of the terminal evolved populations selected without 3-AT (C; n = 76 clones) and with 2.5 mM 3-AT (D; n = 80 clones). Wild-type and WT-synonymous clones are grey. In (C), WT and WT-synonymous frequencies are read against the left axis and individual substitution frequencies against the expanded right axis. Asterisks (e.g., Q12\*, W114\*) denote nonsense (premature-stop) substitutions. **E** Synonymous and non-synonymous mutation frequencies with and without 3-AT. Only non-synonymous mutation counts differed (two-sided Mann-Whitney U test).

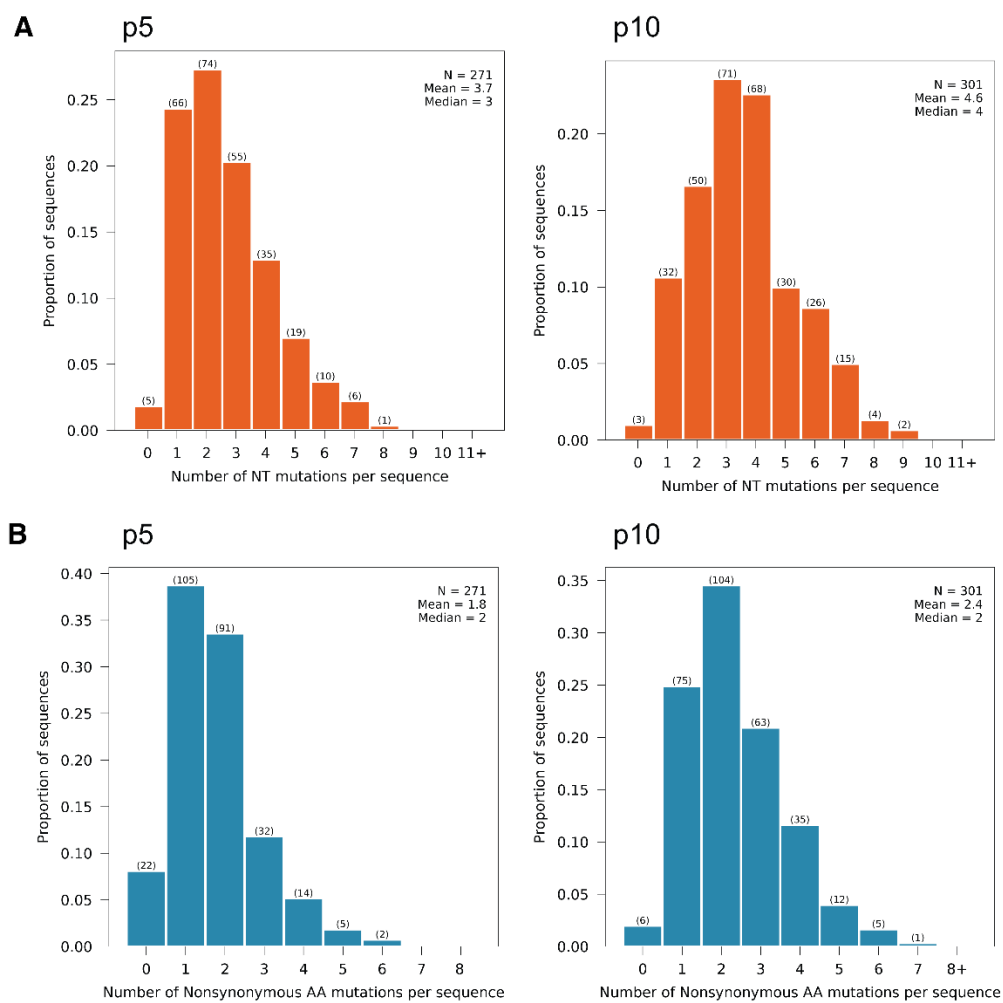

**Supplementary Fig. 20. Mutational frequency of OPRD1 from passage 5 to passage 10.** Distributions of mutations per clone in OPRD1 variants after five (p5) and ten (p10) passages of automated (robot-driven) OrthoRep continuous evolution against 100 μM looperamide, from long-read consensus sequencing of individual clones. **A** Nucleotide (NT) substitutions per sequence at p5 ($n = 271$, mean 3.7, median 3) and p10 ($n = 301$, mean 4.6, median 4). **B** Non-synonymous amino-acid (AA) substitutions per sequence for the same clones at p5 (mean 1.8, median 2) and p10 (mean 2.4, median 2). Bars show the proportion of sequences per bin, with clone counts in parentheses; summary statistics are inset. Both distributions shift toward higher loads with continued passaging.

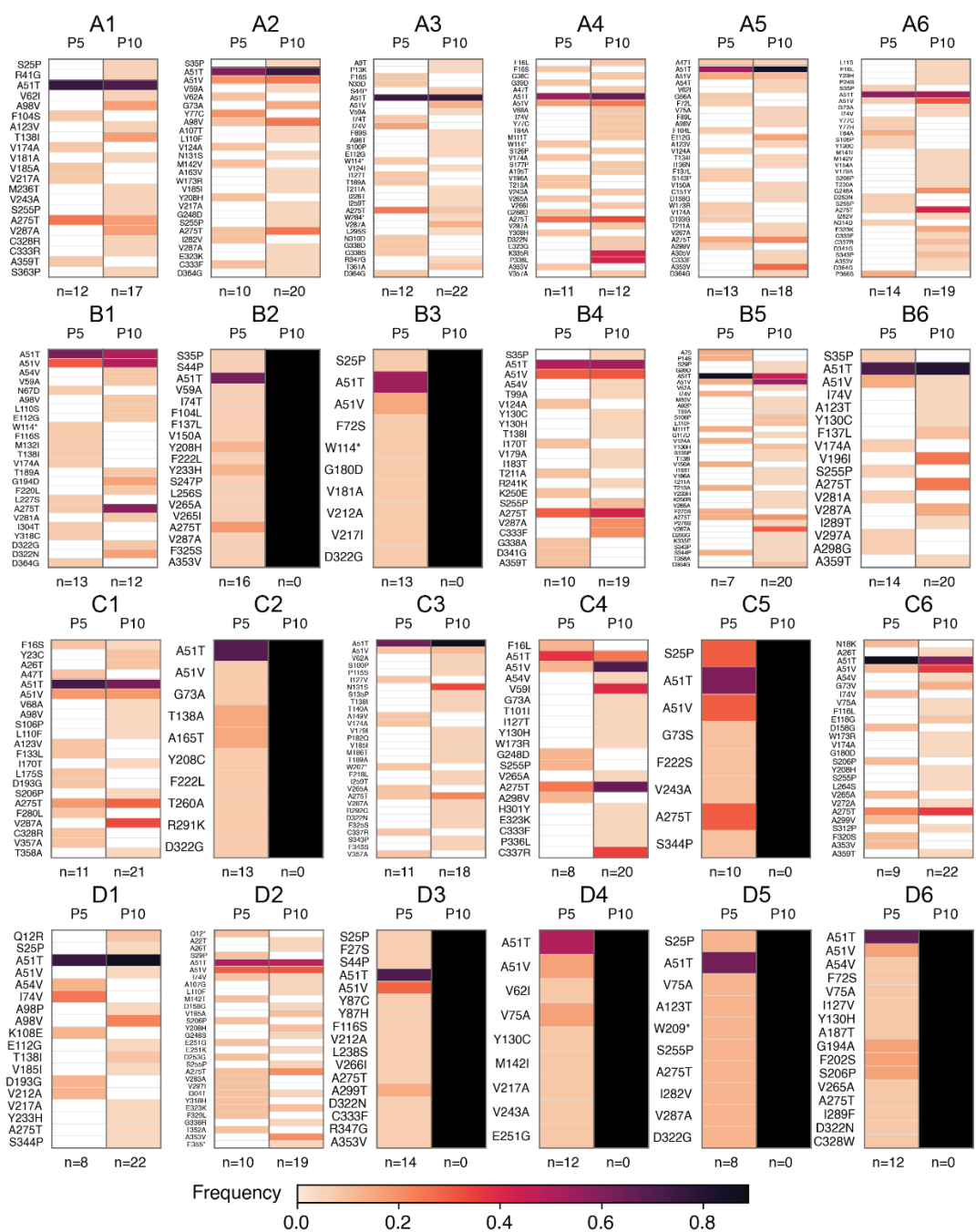

**Supplementary Fig. 21. Well-level resolution of mutational frequency of OPRD1 during automated evolution in the presence of loperamide.** Distributions of OPRD1 substitutions from individual well populations after five (p5) and ten (p10) passages using automated HERO passaging. Shaded frequencies of each substitution are shown for p5 (left) and p10 (right).

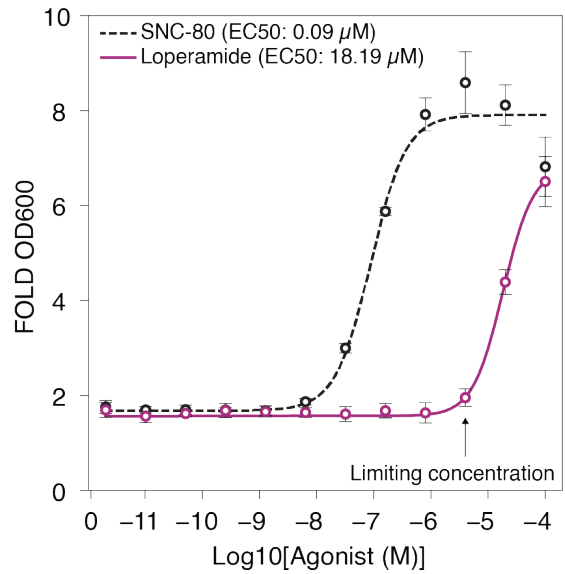

**Supplementary Fig. 22. Single-copy OPRD1 responds to loperamide and SNC-80.** Growth-based dose–response of wild-type OPRD1 integrated as a single copy into the tandem reporter strain, challenged with the δ-opioid agonists SNC-80 (dashed black; $EC_{50} = 0.09 \mu M$) and loperamide (magenta; $EC_{50} = 18.19 \mu M$). Agonist was applied across a serial log-dilution series; response is fold change in $OD_{600}$ relative to the unstimulated control. Points are mean ± s.d. of four replicates. This curve defined the limiting loperamide concentration for OPRD1-dependent growth (arrow); $4 \mu M$ was subsequently selected for high-throughput phenotyping of mutant populations.

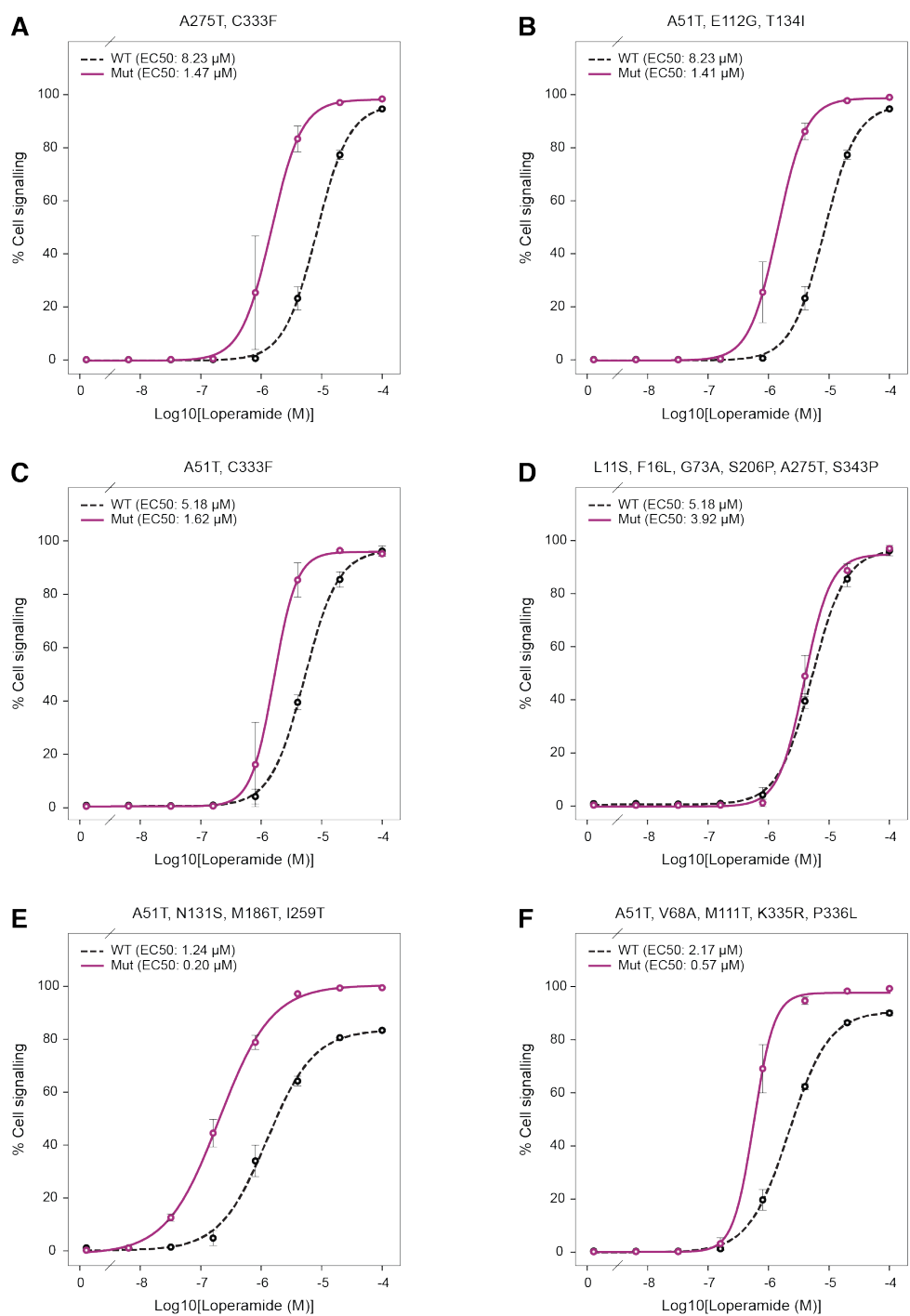

**Supplementary Fig. 23. Additional isolated mutants from OPRD1 high throughput evolution.** Six further receptor variants were picked from the automated high-throughput screen of OPRD1 evolution passage 10 and tested in dose–response format against loperamide. For each variant, cell signaling (% of maximal response) was measured across a loperamide concentration series ($\log_{10}$[loperamide], M) and fit to a sigmoidal dose–response model; the mutant (magenta, solid) is shown against a wild-type reference (black, dashed) assayed in parallel, with fitted EC<sub>50</sub> values inset. WT reference curves were re-run per experimental batch, so absolute EC<sub>50</sub> values should be compared within a panel (mutant vs its paired WT for that run) rather than across panels. Points are means ± s.d. of replicate wells. Each panel is labeled with the non-synonymous substitutions carried by that isolate. Most variants show a pronounced leftward

shift relative to wild type (increased loperamide potency; lower EC<sub>50</sub>); although the shift is modest in **D**. **A** A275T, C333F. **B** A51T, E112G, T134I. **C** A51T, C333F. **D** L11S, F16L, G73A, S206P, A275T, S343P. **E** A51T, N131S, M186T, I259T. **F** A51T, V68A, M111T, K335R, P336L.
